# A genome wide CRISPR/Cas9 screen identifies calreticulin as a selective repressor of ATF6α

**DOI:** 10.1101/2024.02.10.579381

**Authors:** Joanne Tung, Lei Huang, Ginto George, Heather P. Harding, David Ron, Adriana Ordóñez

## Abstract

Activating transcription factor 6 (ATF6) is one of three endoplasmic reticulum (ER) transmembrane stress sensors that mediate the unfolded protein response (UPR). Despite its crucial role in long-term ER stress adaptation, regulation of ATF6 alpha (α) signalling remains poorly understood, possibly because its activation involves ER-to-Golgi and nuclear trafficking. Here, we generated an ATF6α/IRE1 dual UPR reporter CHO-K1 cell line and performed an unbiased genome-wide CRISPR/Cas9 mutagenesis screen to systematically profile genetic factors that specifically contribute to ATF6α signalling in the presence and absence of ER stress. The screen identified both anticipated and new candidate genes that regulate ATF6α activation. Among these, calreticulin (CRT), a key ER luminal chaperone, selectively repressed ATF6α signalling: Cells lacking CRT constitutively activated a BiP::sfGFP ATF6α-dependent reporter, had higher BiP levels and an increased rate of trafficking and processing of ATF6α. Purified CRT interacted with the luminal domain of ATF6α *in vitro* and the two proteins co-immunoprecipitated from cell lysates. CRT depletion exposed a negative feedback loop implicating ATF6α in repressing IRE1 activity basally and overexpression of CRT reversed this repression. Our findings indicate that CRT, beyond its known role as a chaperone, also serves as an ER repressor of ATF6α to selectively regulate one arm of the UPR.

## Introduction

The endoplasmic reticulum (ER), and its associated chaperones, constitutes the major cellular compartment for the synthesis, folding, and quality control of secretory proteins (Sun and Brodsky, 2019). An imbalance between synthesis and folding can lead to ER stress, potentially resulting in ER dysfunction and pathological conditions (Wang and Kaufman, 2016). To restore cellular homeostasis, adaptive pathways, collectively known as the unfolded protein response (UPR), are activated. The UPR is coordinated by three known ER stress transducers: IRE1, PERK and ATF6 (Walter and Ron, 2011). Of the three arms of the UPR, the mechanisms that govern ER stress-dependent ATF6 activation remain the least well understood, even though ATF6 mediates much of the ER-stress-induced changes in gene expression observed in vertebrates. Thus, ATF6 is specialised in the regulation of quality control proteins in the ER (Adachi et al., 2008) and complete loss of ATF6 impairs survival upon ER stress in cellular and animal models (Wu et al., 2007).

ATF6 is a type II transmembrane ER glycoprotein containing an N-terminal portion comprising its basic leucine zipper (b-ZIP), DNA-binding and transactivation domains and a C-terminal ER luminal domain (LD) that senses stress and regulates ATF6α activity. Vertebrates have two isoforms: ATF6α and ATF6β. The ATF6α isoform dominates the regulation of UPR target genes in mammalian cells and is the focus of this study. Unlike IRE1 and PERK, which remain in the ER after UPR activation, ATF6 translocates from the ER to the Golgi, where it is sequentially cleaved by two serine protease, site-1 (S1P) and site-2 (S2P) proteases (Haze et al., 1999) to release its soluble active N-cytosolic domain (N-ATF6α) that is further translocated into the nucleus. There, N-ATF6α acts as a potent but short-live transcription factor (George et al., 2020) that binds ER stress response elements (ERSE-I and -II) in the promoter regions of UPR target genes in complex with the general transcription factor NF-Y and YY1 to upregulate the expression of a large number of ER chaperones (Li et al., 2000; Yoshida et al., 2000). Among those, the gene encoding the ER Hsp70 chaperone BiP (*GRP78*) has been well documented as an important ATF6α target (Adachi et al., 2008).

At ER level, it has been proposed that ATF6α is retained by its association with BiP (Shen et al., 2002), yet the existence of additional factors influencing its ER localisation remains uncertain. Redox regulation has also been proposed to keep ATF6α in the ER as a mixture of monomeric and multimeric disulphide-linked forms (Nadanaka et al., 2007) that upon ER stress are reduced by protein disulphide isomerase (PDI) family members (such as ERp18 or PDIA5), promoting the transit of a reduced monomeric ATF6α population (Higa et al., 2014; Oka et al., 2019). COPII-coated vesicles have been implicated in the trafficking of ATF6α (Schindler and Schekman, 2009; Lynch et al., 2012) and the ATF6α_LD’s N-glycosylation state has also been proposed to influence its ER trafficking (Hong et al., 2004), yet the details of these processed remain incompletely understood. Furthermore, it is widely recognised that a key step of ATF6α activation occurs in the Golgi, the site of proteolytic processing. This is reflected in the inability of cells lacking S2P to trigger BiP transcription in response to ER stress (Ye et al., 2000). Hence, although there is a relatively clear understanding of the events triggering ATF6α cleavage and activation upon reaching the Golgi, much remains unknown about the factors influencing ATF6α regulation in the ER.

ATF6α signalling also involves crosstalk with the IRE1 pathway. This is partly mediated by direct heterodimerisation of N-ATF6α with XBP1s the active, spliced form (Yamamoto et al., 2007, 2004). However, the relationship between the ATF6α and IRE1 pathway is complex: (i) XBP1 and IRE1 genes are transcriptional targets of activated ATF6α (Yoshida et al., 2001), (ii) XBP1s and IRE1 have some different target genes (Shoulders et al., 2013), (iii) and ATF6α activity can also repress IRE1 signalling (Walter et al., 2018).

Here, we established an ATF6α/IRE1 dual UPR reporter cellular model to conduct an unbiased genome-wide CRISPR/Cas9 screen focussed on ATF6α. Complemented by bioinformatic analysis, post-screen hit validation using targeted gene editing and *in vitro* biophysical assays we report on a comprehended analysis of a complex signalling pathway.

## Results

### A dual UPR reporter cell line to monitor ATF6α and IRE1 activity

In vertebrates, the transcriptional activation of BiP is predominantly governed by ATF6α (Adachi et al., 2008; Adamson et al., 2016). To dynamically monitor ATF6α activity in live cells, a “landing pad” cassette, containing recombination sites and expressing the *Cricetulus griseus* (*cg*) BiP promoter region fused with a super folder green fluorescent protein (BiP::sfGFP), was targeted to the *ROSA26* safe-harbour locus of Chinese hamster ovary cells (CHO-K1) using CRISPR/Cas9 (Gaidukov et al., 2018) (Supplemental S1A) in a previously characterised cell line (*CHO-XC45)* in the lab carrying an integrated XBP1s::mCherry reporter transgene monitoring the IRE1 pathway (Harding et al., 2019). Flow cytometry analysis of these ATF6α/IRE1 dual UPR reporter cells (referred as *XC45-6S*) treated with ER stress-inducing agents such as tunicamycin (which inhibits N-glycosylation), 2-Deoxy-D-glucose (which inhibits glycolysis), or thapsigargin (which depletes ER calcium), revealed time-dependent activation of both UPR arms with a broad dynamic range (Fig 1A, top panel, and Supplemental S1B). Additionally, the introduction of the reporter systems did not affect basal BiP protein levels when compared with plain CHO-K1 cells (Supplemental S1C).

**Fig 1.**
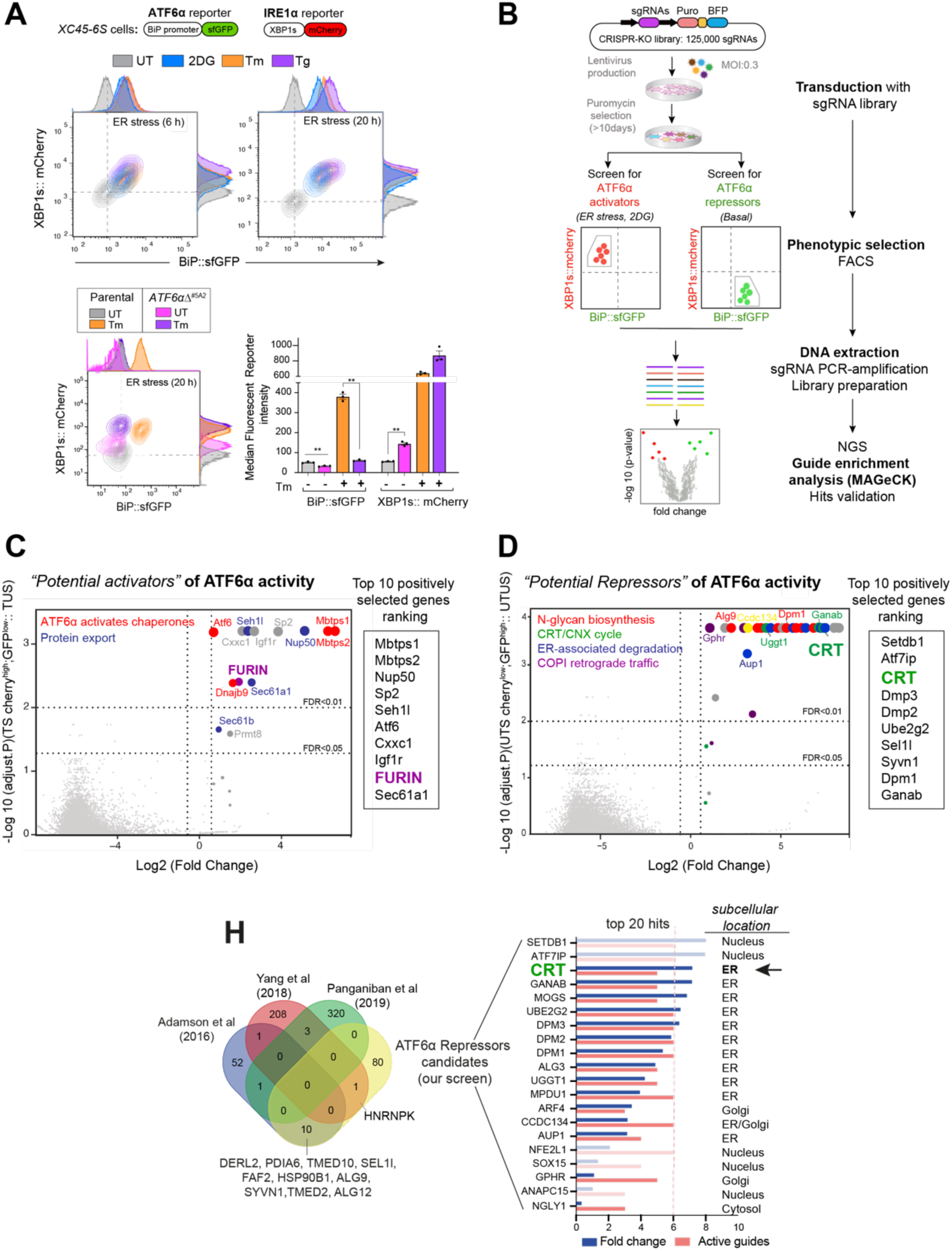
High-throughput CRISPR/Cas9 screens to identify new modulators of ATF6α signalling. **(A)** Characterisation of the *XC45-6S* cells, a dual UPR reporter CHO-K1 cell line stably expressing XBP1s::mCherry to report on IRE1 activity and BiP::sfGFP to report on ATF6α activity that was generated to perform the CRISPR/Cas9 screens. *<u>Top panel:</u>* Two-dimensional contour plots of BiP::sfGFP and XBP1s::mCherry signals in untreated cells (UT, grey) and cells treated with the ER stressors 2-Deoxy-Dglucose (2DG, 4 mM, blue), tunicamycin (Tm, 2.5 µg/ml, orange), or thapsigargin (Tg, 0.5 µM, purple) over a short (6 h) and long period of time (20 h). Histograms of the signal in each channel are displayed on the axes. *<u>Lower panel:</u>* Contour plots as in “1A, top panel” from *XC45-6S* cells (parental) and a knockout *ATF6*αΔ clone (*ATF6*αΔ^#5A2^) generated by CRISPR/Cas9 gene editing. Cells were analysed under basal (UT) and ER stress conditions (Tm, 2.5 µg/ml, 20 h). The bar graph shows the mean ± SEM of the median fluorescent reporter intensity normalised to untreated cells from 3 independent experiments. Statistical analysis was performed by a two-sided unpaired Welch’s t-test and significance is indicated by asterisks (** p < 0.01). **(B)** Flowchart illustrating the two parallel CRISPR/Cas9 screenings aimed to identify genetic ATF6α regulators. ATF6α/IRE dual reporter cells were transduced with a CHO sgRNA CRISPR knockout (KO) library. After puromycin selection, the transduced population was split into two subpopulations, subjected to either ER stress with 2DG treatment (4 mM) for 18 h or left untreated (basal) to identify activators and repressors of ATF6α, respectively. A BiP::sfGFP^low^ ;XBP1s::mCherry^high^ population under ER stress and a BiP::sfGFP^high^; XBP1s::mCherry^low^ population devoid of ER stress were collected through FACS and processed to determinate sgRNAs frequencies after two rounds of selection, expansion, and enrichment. **(C)** Volcano plot depicting the Log2 (fold change) and the negative Log10 (adjusted p value) of the genes targeted by sgRNAs in 2DG-treated-sorted (TS) cells. This analysis identified genes whose loss confers repression of the ATF6α reporter during ER stress without impacting ER-stress-induced IRE1, compared to treated-unsorted (TUS) cells. The table lists the top 10 genes positively selected in the ATF6α *activator screen*, with FURIN highlighted as a hit for further analysis. **(D)** Volcano plot, as in “1C”, depicting the genes targeted by sgRNAs in untreated-sorted (UTS) cells. This analysis identified genes whose loss confers activation of the ATF6α reporter under basal conditions, compared to untreated-unsorted (UTUS) cells. The table lists the top 10 genes positively selected in the ATF6α *repressor screen*, with calreticulin (CRT) highlighted as the hit for further focused analysis. **(H)** Venn-diagram depicting unique and common upregulated top genes examined in “1D”, between our screen (yellow) and previous high-throughput UPR screenings (Adamson et al., 2016; Panganiban et al., 2019; Yang et al., 2018) to discard hits that were confirmed to be IRE1- or PERK-dependent UPR target genes. The Log2 (fold change) and the number of active sgRNA are provided for the 20 genes that remained in our screen, to be put forth as possible ATF6α repressor candidates.

Disruption of endogenous *cgATF6*α in *XC45-6S* cells by CRISPR/Cas9 genome editing abolished the responsiveness of the BiP::sfGFP reporter to ER stress and derepressed IRE1 signalling, both basally and in response to ER stress. These observations are consistent with ATF6α’s obligatory role in BiP expression and with findings that IRE1 signalling is repressed by ATF6α activity (Walter et al., 2018).

To further validate these UPR reporters, cells were subjected to tunicamycin treatment in the presence of Ceapin-A7 (a small molecule that blocks ATF6α activation by tethering it to the lysosome) (Gallagher et al., 2016) or an S1P inhibitor (PF429242) (Hawkins et al., 2008). Ceapin-A7 effectively blocked the activation of the ATF6α fluorescent reporter, whereas the S1P inhibitor partially attenuated the BiP::sfGFP signal in stressed cells (Supplemental S1D).

The selective IRE1 inhibitor 4μ8C (Cross et al., 2012), did not affect the ATF6α-dependent BiP::sfGFP reporter under ER stress conditions, but blocked the IRE1-dependent XBP1s::mCherry reporter (Supplemental S1E). Collectively, these observations confirm that the dual UPR reporter *XC45-6S* cell line enables independent monitoring of ATF6α and IRE1 activities, making it well-suited for identifying additional modulators of ATF6α signalling by high-throughput techniques.

### Genome-wide CRISPR/Cas9 knock-out screens to profile ATF6α signalling

To identify regulators of ATF6α, a library of pooled lentiviral single guide RNAs (sgRNAs) targeting 20,680 predicted protein-coding genes in CHO cells, with ∼ 6 sgRNAs per gene, was used (Ordóñez et al., 2021). Transduction of the *XC45-6S* ATF6α/IRE1 dual UPR reporter cells was performed at a coverage of ∼ 640X, with a multiplicity of infection of 0.3, intentionally set low to disfavour acquisition of more than one sgRNA by any one cell (Shalem et al., 2014). Non-transduced cells were purged by puromycin selection, and the pool of transduced cells was expanded for 10 days to favour gene editing and the establishment of loss of-function phenotypes.

To systematically identify genes regulating the ATF6α transcriptional programme, two positive selection screens were conducted in parallel: a “loss-of-ATF6α function screen” under ER stress conditions to identify activators/enhancers of ATF6α, and a “gain-of-ATF6α function screen” under basal conditions to identify ATF6α repressors (Fig 1B). In the ATF6α *activator screen*, cells were stressed by exposure to 2-Deoxy-D-glucose for 18 h before fluorescent-activated cell sorting (FACS). Cells at the lowest 2% of BiP::sfGFP signal, with preserved IRE1 signalling (BiP::sfGFP^low^; XBP1s::mCherry^high^), were selected for further analysis. The search for genes whose inactivation de-represses ATF6α (ATF6α *repressor screen*) was conducted in the absence of ER stress. Here, cells exhibiting a constitutively active ATF6α reporter but only basal levels of the IRE1 reporter (BiP::sfGFP^high^; XBP1s::mCherry^low^) were selected for further analysis. Sorted cells from both screens underwent two rounds of progressive enrichment through cell culture expansion and phenotypic selection by sorting. As a result, a progressive increase in the proportion of BiP:sfGFP^low^ or BiP:sfGFP^high^ cells occurred in the respective screen (Supplemental S2A and S2B).

The integrated sgRNAs were amplified from genomic DNA, and their abundance in the enriched populations after one or two rounds of sorting was estimated by deep sequencing and compared to a reference sample of transduced and unsorted population. MAGeCK bioinformatics analysis (Li et al., 2014) was used to determine sgRNA sequence enrichment, establishing the corresponding ranking of targeted genes (Table S1; see GEO accession number: GSE254745) and quality controls (QC) measures (Supplemental S2C and S2D). Both screens were conducted at least in duplicate for all conditions.

### Expected and new candidate genes that contribute to ATF6α activation

MAGeCK-based analysis of sgRNAs enriched in cells with attenuated ATF6α signalling and intact activity of the IRE1 reporter, revealed that those targeting S1P and S2P proteases (encoding by the *MBTPS1* and *MBTPS2* genes) were the two most enriched genes (Fig 1C), validating the experimental methodology. Gene Ontology (GO) cluster analysis of the 100 most significantly enriched targeted genes revealed up to 19 enriched pathways (Supplemental S3A), with the top two pathways highlighted in Figure 1C being closely related to ATF6α signalling.

We focused our attention on targeted genes that could act upstream to find early regulatory events of ATF6α. Therefore, targeted genes that were likely to act downstream and impact on the cleaved ATF6α form (N-ATF6α) were dismissed, including the *SP2* transcription factor, which shares the binding site with NF-Y (Völkel et al., 2015), and also genes involved in nuclear import, such as *NUP50* and *SEH1L* (Fig 1C and Supplemental S3A). Similar consideration was applied to *PRMT8*, as the family member *PRMT1* has been shown to enhance ATF6α transcriptional activity (Baumeister et al., 2005). Additionally, guides targeting *IGFR1* and *CXXC1* were also dismissed as likely affecting ER proteostasis indirectly *via* cellular growth (Pfaffenbach et al., 2012). Guides targeting the genes encoding the ER resident proteins *SEC61A1*, *SEC61B* (both components of the translocon machinery) and the *DNAJB9*/ERDj4 co-chaperone were highly enriched. However, given their roles in selectively regulating IRE1 signalling (Adamson et al., 2016; Shoulders et al., 2013), we deemed their identification to reflect a bias towards IRE1 activators arising from the selection scheme for ATF6α^low^ and IRE1^high^ cells.

One of the genes that could plausibly play a proximal role and act upstream on ATF6α activation was the ubiquitous Golgi-localised protease FURIN (Figure 1C). Like S1P, FURIN belongs to the subtilisin-like proprotein convertase family (Nakayama, 1997; Van de Ven et al., 1991) and is thus poised to act at the level of ATF6α proteolytic processing. All six sgRNAs targeting *FURIN* were enriched in the BiP:sfGFP^low^ cells, a level of enrichment similar to that observed to the established ATF6α regulators *MBTPS1* (S1P) and *MBTPS2* (S2P) (Supplemental S3B). The genotype-phenotype relationship suggested by the screen was also observed upon targeting *FURIN* in ATF6α/IRE1 dual reporter cells, as both pools of targeted cells and an individual clone had diminished responsiveness of BiP:sfGFP to ER stress (Supplemental S4A left and middle panels). Furthermore, the S1P inhibitor more effectively attenuated the BiP::sfGFP reporter in *FURIN*Δ cells compared to wild-type cells (Supplemental S4A right panel, S4B and S4C).

These observations raised the possibility of redundant roles of FURIN and S1P in activating ATF6α by cleaving its luminal domain. However, ATF6α_LD expressed and purified from mammalian cells, failed to serve as a substrate for FURIN *in vitro* (Supplemental S4D), suggesting instead that FURIN’s role in regulating ATF6α signalling is indirect.

### Calreticulin and an interconnected gene network repress ATF6α

To identify genes that basally repress ATF6α, a similar MAGeCK-based data analysis was applied to the targeted genes enriched in cells that selectively activated the ATF6α reporter under basal conditions. GO analysis of the 100 most significantly enriched genes established an interconnected network highlighting signalling processes including the calreticulin (CRT)/calnexin (CNX) cycle, N-glycan biosynthesis, ERAD and COPI-dependent Golgi-to ER retrograde traffic (Fig 1D and Supplemental S3C). Among these, the top 10 genes included CRT, components of the dolichol-phosphate mannose (DPM) synthase complex, such as *DPM3*, *DPM2* and *DPM1* involved in the N-glycosylation process, and ERAD components, such as *UBE2G2, SEL1L* (Fig 1D).

Because these four pathways potentially activate all three branches of the UPR, genes previously identified in genome-wide screens to modulate IRE1 or PERK functionality were excluded from further analysis (e.g. *TMED10, SEL1L, FAB2, SYVN1, PDIA6*) (Fig 1H) (Adamson et al., 2016; Panganiban et al., 2019; Yang et al., 2018). Of the remaining genes, we focused on those encoding proteins localised in the ER lumen or Golgi, which could act as specific proximal regulators of ATF6α. Among these, CRT, an abundant luminal ER lectin chaperone, caught our attention (Fig 1H). Notably, although CRT promotes the folding of ER synthesised glycoproteins *via* the CRT/CNX cycle (Lamriben et al., 2016), its ER transmembrane homolog CNX was not identified in the screen. This observation was further supported by the fact that sgRNAs targeting *CNX* were not enriched, whereas five out of six sgRNAs targeting *CRT* showed significant enrichment during the selection process (Fig 2A).

**Fig 2.**
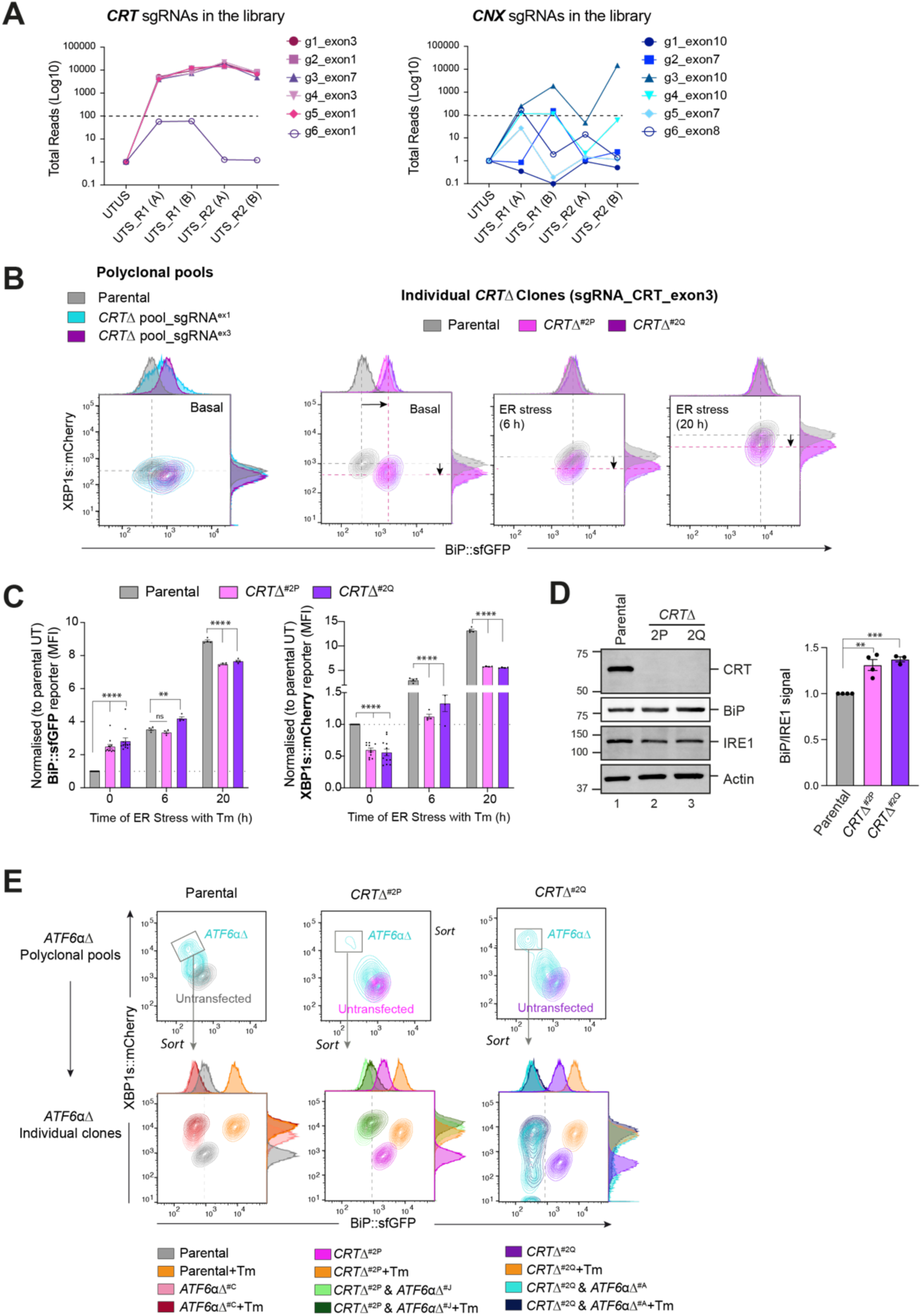
Calreticulin-depleted cells exhibit constitutive activation of ATF6α signalling. **(A)** Total read count enrichment through the selection process for each active sgRNA targeting calreticulin (CRT) and calnexin (CNX). [UTUS: untreated and unsorted; UTS: Untreated and sorted; R1: Round 1 of enrichment; R2: Round 2 of enrichment; A-C: pools of cells; g1-g6: sgRNAs]. **(B)** Two-dimensional contour plots of BiP::sfGFP and XBP1s::mCherry signals examined in two derivative *CRT*Δ polyclonal pools (left) and two independent *CRT*Δ clones (right, named *CRT*Δ^#2P^ and *CRT*Δ^#2Q^) under basal conditions or ER stress induced with 2.5 µg/ml Tm for a short (6 h) and extended period of time (20 h). A representative dataset from more than four independent experiments is shown. **(C)** Normalised quantification expressed as fold-change of the median fluorescence intensity (MFI) of BiP::sfGFP (left) and XBP1s::mCherry signals (right) in the two independent *CRT*Δ clones from more than four independent experiments, as described in “2B”, indicated by mean ± SEM. **(D)** Representative immunoblots of endogenous CRT and BiP protein levels in cell lysates from *XC45-6S* parental cells and two *CRT*Δ derivatives clones selected for functional experiments. The samples were also blotted for IRE1 and actin (loading controls). The right graph bar shows the ratio of BiP to IRE1 signal in four independent experiments indicated by mean ± SEM. **(E)** *<u>Top panel.</u>* Two-dimensional contour plots of BiP::sfGFP and XBP1s::mCherry signals in polyclonal pools of parental cells and two *CRT*Δ clones after depleting the endogenous *ATF6*α locus by CRISPR/Cas9 gene editing. *<u>Lower panel.</u>* Grey rectangles in top panels indicate the subcellular polyclonal population re-sorted to analyse single *ATF6*αΔ clones in parental and *CRT*Δ clones. ER stress treatments with 2.5 µg/ml Tm lasted 20 h. All statistical analysis was performed by two-sided unpaired Welch’s t-test and significance is indicated by asterisks (** p<0.01; *** p < 0.001; **** p < 0.0001).

### Constitutive activation of ATF6α in cells lacking calreticulin

To explore *CRT*’s role in ATF6α regulation, *CRT* was targeted by CRISPR/Cas9 in wild-type *XC45-6S* dual UPR reporter cells. *CRT*-targeted cells exhibited large populations of BiP::sfGFP^high^; XBP1s::mCherry^low^ cells, mirroring the results of the initial screen. This phenotype was stable in single clones from the targeted pool (Fig 2B and 2C). Targeting CRT repressed the IRE1 reporter basally and left little room for further induction of the ATF6α reporter upon ER stress (Fig 2B and 2C). De-repression of the ATF6α reporter in *CRT*Δ clones correlated with a slight increase in BiP protein levels (Fig 2D). Immunoblots of ER-resident proteins (reactive with antibody directed to their endogenous KDEL C-terminal tag) confirmed the increase in BiP protein levels in *CRT*Δ cells and revealed no marked changes in GRP94 protein levels, a slight increase in P3H1 levels, and a notably increase in a band likely corresponding to PDIA in a *CRT*Δ clone (Supplemental S5). By contrast, depletion of CNX had no impact on the BiP::sfGFP reporter (Supplemental S6A-S6C), consistent with the findings of the screen.

To test whether de-repression of the BiP::sfGFP reporter in *CRT*Δ cells was ATF6α dependent, endogenous *ATF6*α was also inactivated in *CRT*Δ cells. ATF6α inactivation restored BiP::sfGFP reporter to its low baseline levels and derepressed the IRE1 reporter to levels observed in *XC45-6S* parental cells (Fig 2E). Overall, these findings confirmed that BiP::sfGFP reporter activity correlates with the transcriptional activity of ATF6α in *CRT*Δ cells.

### Calreticulin represses Golgi-trafficking and processing of ATF6α

Considering the crucial role of trafficking in ATF6α activation, we next investigated whether depletion of CRT affected the stress-induced localisation of ATF6α by comparing the subcellular localisation of ATF6α in parental and *CRT*Δ cells. As trafficking is closely linked to proteolytic processing, wild-type ATF6α does not accumulate in the Golgi of stressed cells (Shen et al., 2002). To stabilise ATF6α molecules reaching the Golgi, a CHO-K1 cell line stably expressing a GFP-tagged version of *cg*ATF6α_LD lacking S1P and S2P cleavage sites (GFP-ATF6α_LD^S1P,S2Pmut^) was engineered (Fig 3A). In non-stressed cells, GFPATF6α_LD^S1P,S2Pmut^ was distributed in a prominent reticular ER pattern with only a minor fraction co-localising with the Scarlet-tagged Giantin Golgi localisation marker. Exposure to the rapidly acting ER stress-inducing agent Dithiothreitol (DTT), increased the Golgi pool at the expense of the ER pool (Fig 3A); an expected outcome validating GFP-ATF6α_LD^S1P,S2Pmut^ as a sentinel for the ATF6α protein. Compared to parental cells, GFP-ATF6α_LD^S1P,S2Pmut^ showed significantly more Golgi co-localisation in *CRT*Δ cells under basal conditions (Fig 3A). These results were in line with our previous flow cytometry observations and suggested that CRT contributes to ER retention of ATF6α, either directly or indirectly.

**Fig 3.**
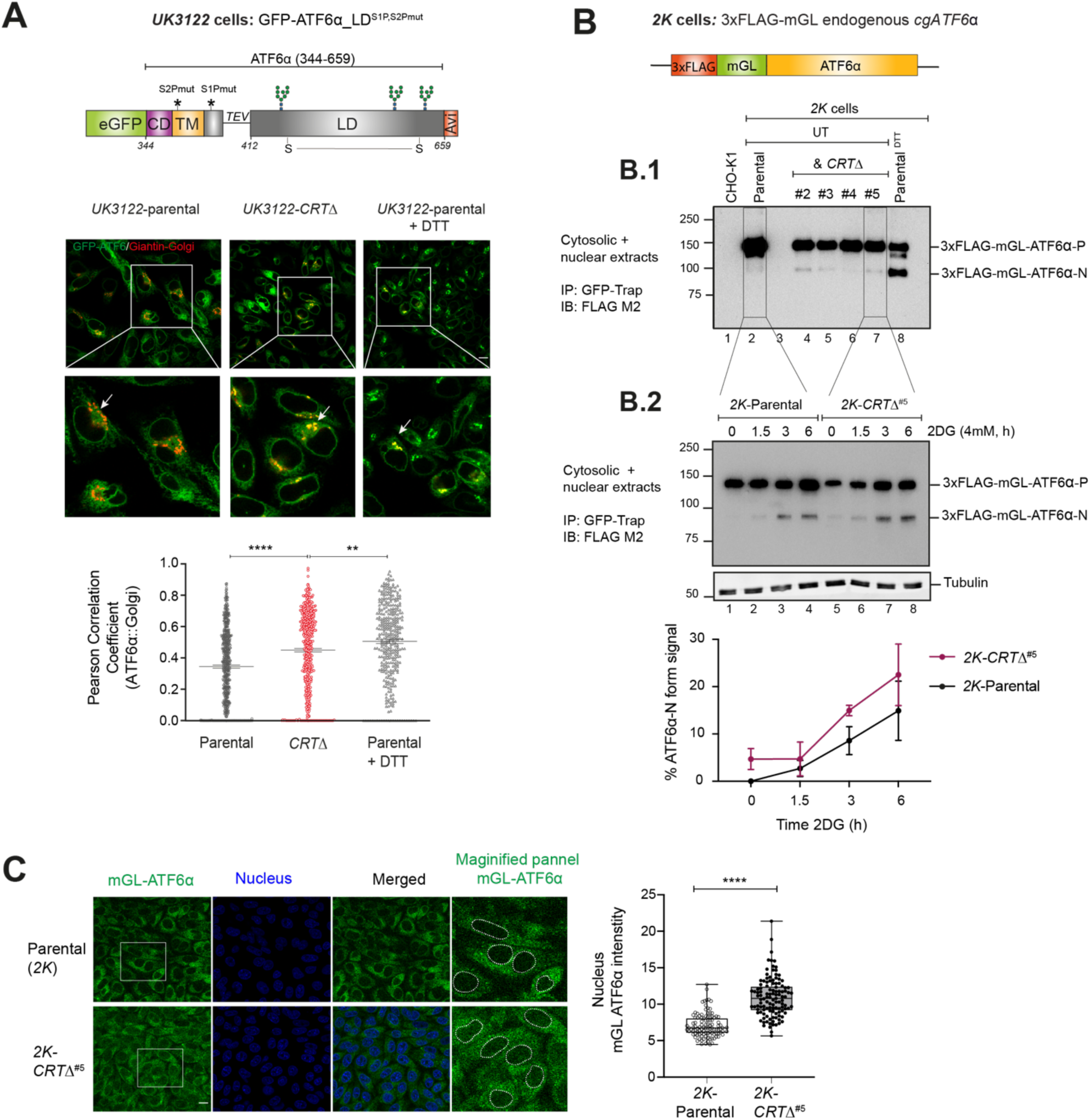
Loss of calreticulin promotes higher proportion of ATF6α trafficking to Golgi and nucleus. **(A)** *<u>Top panel:</u>* Schematic representation of the genetic construct (UK3122) designed for generating a CHO-K1 stable cell line expressing a N-terminus tagged ATF6α, where eGFP replaces most of the cytosolic domain of *cg*ATF6α, and the S2P and S1P cleavage sites are mutated to increase the intact fraction in the Golgi. Colour code: eGFP tag in green; cytosolic domain (CD) in purple; transmembrane domain (TM) in yellow; TEV protease cleavage site is represented with a linker line; luminal domain (LD) in grey; and a C-terminus Avi-tag in red. *<u>Lower panels:</u>* Representative live cell confocal microscopy images of *UK3122* CHO-K1 cells stably expressing GFP-ATF6α_LD^S1P,S2Pmut^ and transiently expressing a pmScarlet _Giantin-C1 plasmid as a red fluorescent Golgi marker. Both parental cells and a CRISPR/Cas9 gene edited *CRT*Δ derivative pool were imaged. DTT treatment (4mM, 1h) was applied to parental cells to positively influence ATF6α’s Golgi localisation. Each panel’s magnified sections are presented in the insets. Arrows indicate GFP-ATF6α_LD^S1P,S2Pmut^::Golgi co-localisation. Pearson coefficients for the co-localisation of GFP-ATF6α_LD^S1P,S2Pmut^with the Golgi apparatus marker Giantin in parental cells (n >150), parental cells treated with 4 mM DTT for 1 h (n >100) and *CRT*Δ cells (n >150) are presented in the bar graph, including cells from three independent experiments. Volocity software was used for co-localisation quantification. Statistical analysis was performed by a two-sided unpaired Welch’s t-test and significance is indicated by asterisks (** p<0.01; **** p < 0.0001). **(B)** Schematic representation of engineered stable *2K* cells with the endogenous *cgATF6*α locus (in orange) tagged with 3xFLAG (in red) and mGreenLantern (mGL) (in green) at the N-terminus. (**B.1)** Cell lysates from *2K* cells and four independent *2K*-*CRT*Δ derivative clones were harvested, immunoprecipitated (IP) using GFP-Trap Agarose (ChromoTek) and analysed by immunoblot (IB) using an anti-FLAG M2 antibody to detect processing of ATF6α. Full-length ATF6α-Precursor (ATF6α-P) and processed ATF6α-N forms were identified. Treatment with DTT for 1 h (4 mM) was used to induce ATF6α processing. Parental untagged cells were used as control for background. Data shown is representative of one experiment. **(B.2)** Parental *2K* cells and a derivative single *CRT*Δ clone (#5) were treated with 4mM 2DG at the indicated time points (1.5, 3 and 6 h). Cells were harvested and analysed by immunoblot as in “B.1”. Lower panel shows the percentage of ATF6α-N form signal/total ATF6α signal from three independent experiments indicated by mean ± SEM. **(C)** ATF6α nuclear translocation live cell microscopy assay showing unstressed *2K* parental cells and a *2K*-*CRT*Δ clone. mGL-ATF6α is in green, and nucleus in blue. mGL-ATF6α signal intensity in the nucleus (shown by dashed lines) was measured using Volocity software. Data from parental cells (n >100) and *CRT*Δ cells (n > 100) are displayed in a box and whiskers graph, displaying all points (with min. and max. intensities).

GFP-ATF6α_LD^S1P,S2Pmut^ reporter cannot be proteolytically processed. Consequently, to gauge the effect of CRT on ATF6α processing we sought to create a CHO-K1 stable cell line to track endogenous ATF6α. For that, the *cgATF6*α locus was tagged with a 3xFLAG and a monomeric GreenLantern (mGL) fluorescent protein, generating a cell line referred to as *2K*: 3xFLAG-mGL-ATF6α (Fig 3B). The functionality of this probe was confirmed by treatment with DTT that resulted in the appearance of a prominent faster migration band in SDS-PAGE, compatible with the processed S2P-cleaved ATF6α (N-ATF6α) form, and depletion of the precursor form (ATF6α-P) (Fig 3B.1, lane 8). Subsequently, *2K* cells were targeted to generate *2K-CRT*Δ derivatives cells, enabling a comparison of ATF6α processing. *2K*-*CRT*Δ clonal cells exhibited elevated levels of the processed N-ATF6α form under both basal and stress-induced conditions (Figure 3B.1 and 3B.2) that correlated with a higher nuclear mGL-ATF6α signal observed by live cell microscopy (Fig 3C). These findings indicated that CRT represses both ER-to-Golgi trafficking and processing of ATF6α into its active form.

It has been reported that ATF6α can exist in three redox forms: a monomer and two inter-chain disulfide-stabilised dimers (Nadanaka, et al., 2007). To investigate whether CRT depletion alters the redox status of ATF6α, cell lysates from *2K*-parental cells and a *2K*-*CRT*Δ clonal cells were analysed under both basal and ER-stress conditions using non-reducing SDS-PAGE (Supplemental S7A). We detected two redox forms of ATF6α, consistent with an inter-chain disulfide-stabilised dimer and a monomer. Under basal conditions, ATF6α predominantly existed as a high mobility monomer in both *2K*-parental and *2K*-*CRT*Δ cells, with the appearance of an even faster-migrating processed N-ATF6α form in *2K*-*CRT*Δ cells, consistent with the findings presented in Figure 3B. ER stress (induced by 2-deoxy glucose, 2DG) was associated with appearance of a low mobility band consistent with a disulfide-stabilised dimer, and a concomitant decrease in the ATF6α monomer intensity in *2K*-parental cells, as previously reported (Oka et al., 2022). Interestingly, under stress conditions, the monomer levels in *2K*-*CRT*Δ cells remained largely unchanged, indicating a higher monomer-to-dimer ratio compared to parental cells (Supplemental S7B). These data suggest that the loss of CRT may stabilise the monomeric form of ATF6α, which is proposed to be more efficiently trafficked. This observation aligns with our results showing that CRT depletion is linked to activation of ATF6α.

### Calreticulin interacts with the ATF6α luminal domain *in vitro*

To investigate the underlying mechanism of CRT’s repressive effect on ATF6α, we explored the physical interaction between the two purified proteins *in vitro* using BioLayer Interferometry (BLI). The biotinylated ATF6α_LD, expressed in mammalian cells, was immobilised on the BLI probe and used as the ligand, and bacterially expressed CRT was used as the analyte. Considering possible regulatory roles of N-glycosylation and disulphide bonds in ATF6α activation (Hong et al., 2004; Oka et al., 2019), three different versions of ATF6α_LD were used: 1) fully glycosylated ATF6α_LD (WT), 2) non-glycosylated ATF6α_LD (ΔGly) and 3) ATF6α_LD lacking its cysteines (ΔC) (Fig 4A). BLI showed reversible binding of CRT to ATF6α_LD^WT^ that was characterised by slow association (*k*_on_) and dissociation (*k*_off_) rates (Fig 4B). Both the association and dissociation phases were fitted to a single exponential association-dissociation model, yielding a *K*_D_ within the range of 1 µM, a concentration that was consisted with the estimated CRT concentration in the ER of CHOK1 cells obtained through quantitative immunoblotting (∼7 µM) (Supplemental Fig S6D). Similar interactions were also observed with ATF6α_LD^ΔGly^ and ATF6α_LD^ΔC^.

**Fig 4.**
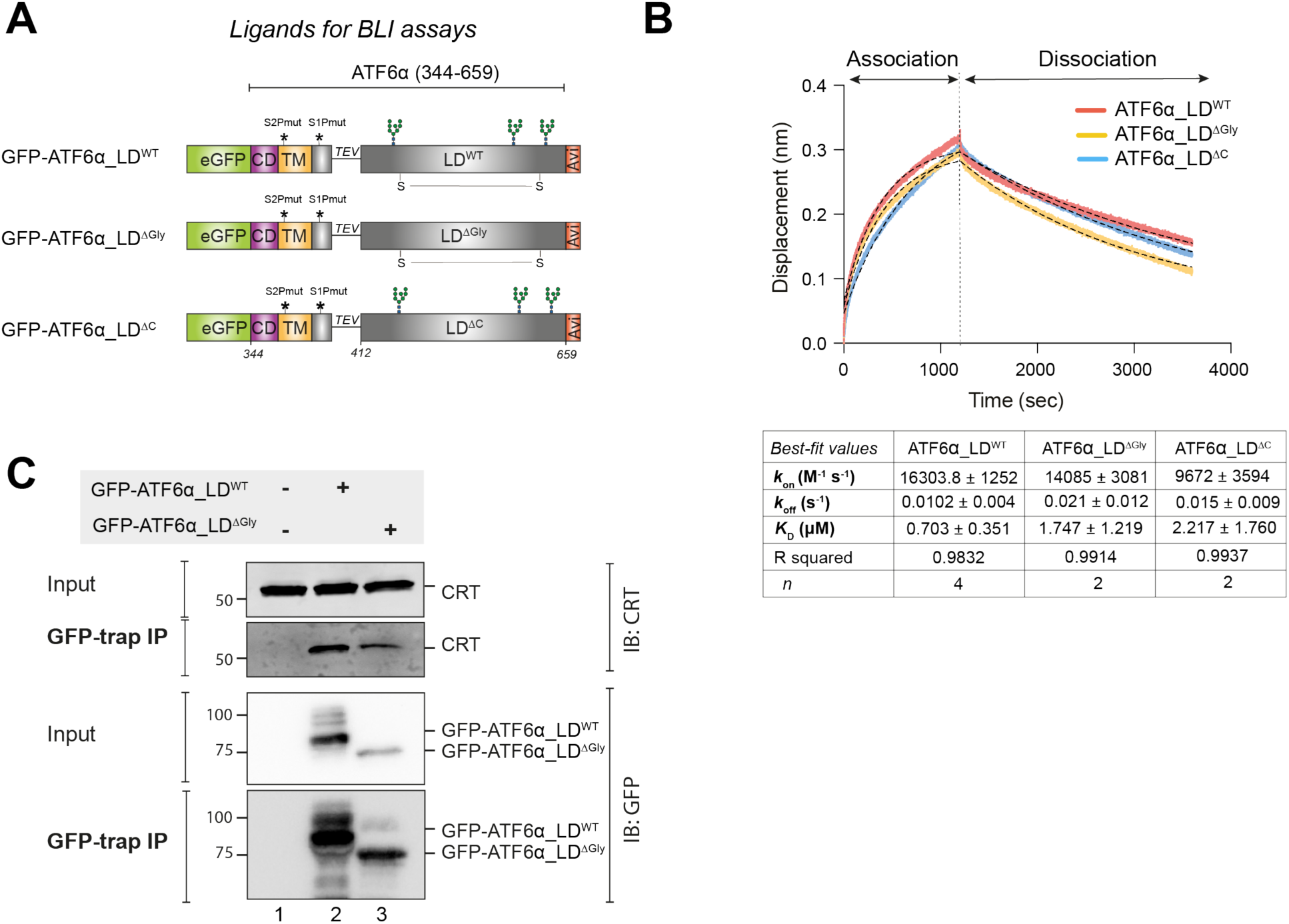
Calreticulin interacts directly with the ATF6α luminal domain *in vitro* and in cells. **(A)** Schematic representation of the constructs designed to produce three luminal domains (LD) of ATF6α to be used as ligands in bio-layer interferometry (BLI) assays: wild-type (WT) ATF6α_LD, nonglycosylated (ΔGly) ATF6α_LD, and ATF6α_LD lacking its cysteines (ΔC). The probes underwent *in vivo* biotinylation, purification using GFP-Trap Agarose (ChromoTek), and were eluted *via* TEV cleavage. Colour code: eGFP tag in green; cytosolic domain (CD) in purple; transmembrane domain (TM) in yellow; TEV cleavage site with a linker line; LD in grey; C-terminus Avi-tag in red. S-S indicates for the disulphide bond in the LD between two cysteines. **(B)** Representative BLI signals depicting the interaction of rat (r) CRT (analyte) with different forms of *cg*ATF6α_LD (from HEK293T cells) immobilised on streptavidin biosensors: ATF6α_LD^WT^ in red, ATF6α_LD^ΔGly^ in yellow, and ATF6α_LD^ΔC^ in blue. The interaction kinetics were calculated over the entire 1200 seconds of the association phase and 2400 seconds of the dissociation phase. Dotted lines represent the best-fit association then dissociation curves using a nonlinear regression to calculate kinetic constants by Prism10. The table represents the BLI measurements of the kinetics of the interaction between rCRT and *cg*ATF6α_LD from 2 - 4 independent experiments (means ± SEM). Interaction kinetics were measured at a rCRT concentration of 10 µM. **(C)** Representative immunoblots (IB) of endogenous CRT recovered in complex with ATF6α in HEK293T cells overexpressing GFP- *cg*ATF6α_LD^WT^, GFP-*cg*ATF6α_LD^ΔGly^ or neither. Immunoprecipitations (IP) were performed using GFP-Trap Agarose, and then membranes were blotted with anti-human CRT and anti-GFP antibodies. The immunoblots are representative for three independent experiments.

To examine a possible interaction between CRT and ATF6α in cells, HEK293T cells were transfected with either GFP-ATF6α_LD^WT^ or GFP-ATF6α_LD^ΔGly^, followed by selective recovery using GFP-Trap Agarose and subsequent immunoblotting with an anti-CRT antibody. The co-immunoprecipitation results showed that both ATF6α_LD^WT^ and, to a lesser extent, ATF6α_LD^ΔGly^ immunoprecipitated endogenous CRT (Fig 4C). Re-probing the blot with an anti-GFP antibody revealed that GFP-ATF6_LD^ΔGly^ was expressed at lower levels than ATF6α_LD^WT^ (Fig 4C), potentially accounting for the reduced recovery of CRT in complex with ATF6α_LD^ΔGly^. Overall, these observations suggested that CRT binds to ATF6α_LD both in cells and in isolation *in vitro*, supporting the notion of a direct interaction between the two proteins.

### A genetic platform to study endogenous ATF6α signalling

To circumvent the potentially corrupting effect of protein overexpression, we sought to express ATF6α variants from the endogenous locus, by replacing the ATF6α_LD coding sequence with mutants of our design. To enable CRISPR-Cas9 mediated homology-directed repair (HDR) that replaces the wild-type sequence with the mutants, we first created a non-functional ATF6α allele that could be subsequently restored to function by homologous recombination, offering wild-type or mutant repair templates (Fig 5A). As expected, the deletion of ATF6α resulted in the loss of the BiP::sfGFP signal in stressed cells (and enhanced XBP1s::mCherry signal in basal conditions, Fig 5A). This ATF6αΔ clone was re-targeted with a unique sgRNA/Cas9, alongside ATF6α_LD^WT^, ATF6α_LD^ΔGly^ or ATF6α_LD^ΔC^ repair templates (Fig 5B).

**Fig 5.**
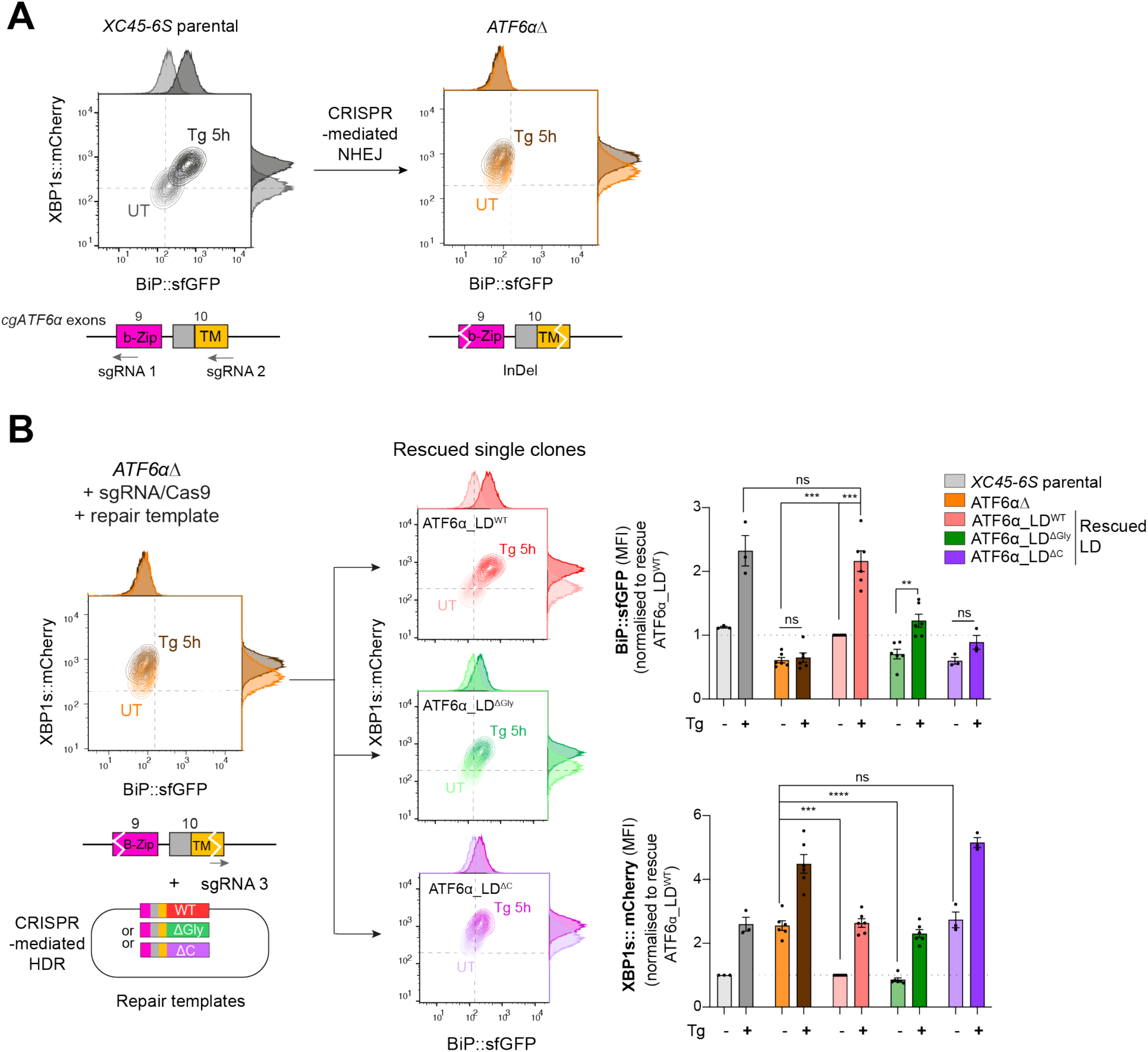
A homologous CRISPR/Cas9 gene editing recombination platform to study ATF6α variants at endogenous levels in CHO-K1 cells. **(A)** Two-dimensional contour plots of BiP::sfGFP and XBP1s::mCherry signals in *XC45-6S* parental cells (in grey; left) and in a null functional ATF6α clonal cell (*ATF6α*Δ, in orange; right) generated by CRISPR/Cas9-mediated NHEJ (non-homologous end joining) to introduce frameshifting mutations into the DNA-binding (b-Zip, pink) and transmembrane (TM, yellow)encoding regions by targeting exons 9 and 10 of the endogenous locus of *cgATF6*α. Cells were analysed in basal conditions (UT) or upon ER induction with Tg (0.5 µM, 5 h). A schema for the sgRNAs/Cas9 that target cgATF6 is shown below the plots. **(B)** *ATF6α*Δ cells were re-targeted with a sgRNA/Cas9 directed to the mutated exon 10 and offering three repair templates: ATF6α_LD^WT^ in red, ATF6α_LD^ΔGly^ in green and ATF6α_LD^ΔC^ in purple. Successfully rescued single clones by CRISPR/Cas9-mediated homology-directed repair (HDR) were evaluated in the absence (pale colour) and presence of ER stress (0.5 µM Tg, 5 h, dark colour). Contour plots are representative from 3-6 independent experiments. Bar graphs display the quantification of the fold-change of BiP::sfGFP (top) and XBP1s::mCherry signals (bottom) in more than three independent experiments indicated by mean ± SEM. Statistical analysis was performed by a two-sided unpaired Welch’s t test and significance is indicated by asterisks (*** p < 0.001; **** p < 0.0001; ns: nonsignificant).

Unlike the ATF6α_LD^WT^, neither the ATF6α_LD^ΔGly^ nor the ATF6α_LD^ΔC^ repair templates fully restored BiP::sfGFP responsiveness to stress (Fig 5B top panel), consistent with the importance of N-glycosylation and disulphide bond formation to ATF6α functionality. Notably, whereas ATF6α_LD^ΔC^ knock-in cells exhibited basal activation of the IRE1 reporter comparable to levels observed in the ATF6αΔ parent (Fig 5B lower panel), the IRE1 reporter signal in the ATF6α_LD^ΔGly^ knock-in cells shifted to that observed in wild-type cells, suggesting that ΔGly allele retained some functionality. The partial ATF6α functionality of the ATF6α_LD^ΔGly^ knock-in allele aligned with the lower expression levels of ATF6α_LD^ΔGly^ compared to ATF6αLD^WT^ in transfected cells, as previously observed in Figure 4C. However, this feature complicated interpretation of experiments designed to establishing the role of ATF6α_LD glycans on CRT-mediated repression of ATF6α: failure of CRT deletion to derepress the BiP::sfGFP signal in ATF6α_LD^ΔGly^ knock-in cells (Supplemental Fig S8) could be interpreted as either an obligatory role for glycans in the repression or as lack of dynamic range of the signal needed to detect a residual effect of CRT deletion in ATF6α_LD^ΔGly^ knockin cells.

### Impact of calreticulin variants on ATF6α reporter activity

CRT is a lectin chaperone that interacts with the protein disulphide isomerase ERp57 to stabilise CRT-substrate binding (Oliver et al., 1999). Therefore, we tested the ability of human CRT^WT^ and two mutants that compromised glycan binding (Y92A) or Erp57 co-chaperone binding (W244A) (Del Cid et al., 2010) to reverse the *CRT*Δ phenotype. Parental and *CRT*Δ clones were transfected with CRT^WT^, CRT^Y92A^ and the double mutant CRT^Y92A,W244A^expressing plasmids, and cells were evaluated by flow cytometry 48 and 72-h post-transfection (Fig 6A). Expression of CRT^WT^ restored basal levels of both BiP::sfGFP and XBP1s:mCherry reporters in *CRT*Δ cells (Fig 6A purple panels, and Fig 6B), confirming the repressive role of CRT on ATF6α. By contrast, expression of the double CRT^Y92A,W244A^ mutant failed to rescue the *CRT*Δ phenotype (Fig 6A green panels, and Fig 6B), whereas CRT*^Y92A^*, selectively incapable of glycan binding, retained some ability to repress BiP::sfGFP and modestly rescuing the *CRT*Δ phenotype (Fig 6A orange panels, and Fig 6B). Immunoblots targeting CRT raised concerns about the integrity of theCRT^Y92A,W244A^ protein, precluding a separate analysis of Erp57’s role in ATF6α repression (Fig 6C). However, CRT^WT^ and CRT^Y92A^ proteins were expressed similarly (Fig 6C). Together, our observations pointed to an important role for interactions between CRT and ATF6α_LD glycans but also left room for some lectin independent repression of ATF6α by CRT.

**Fig 6.**
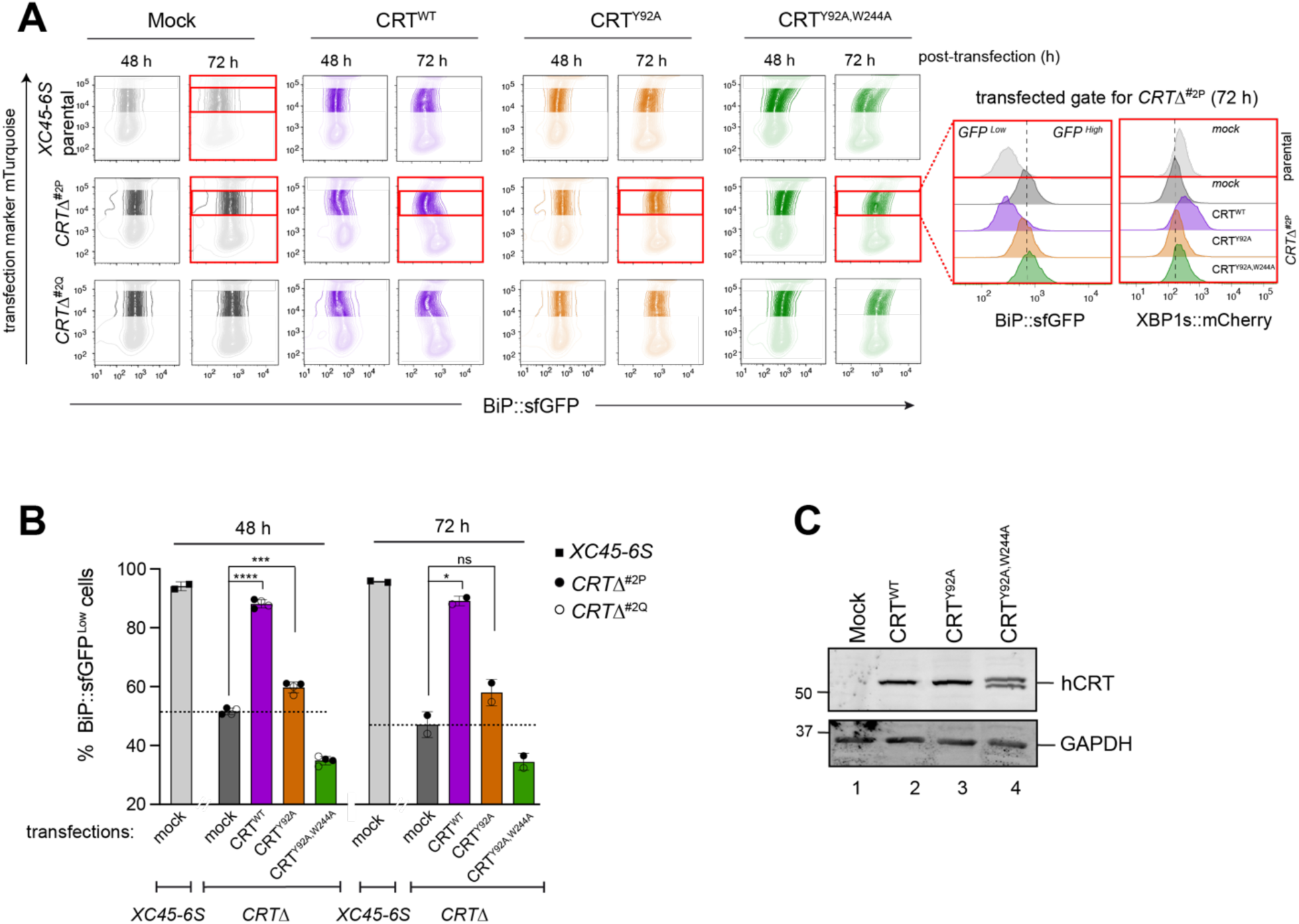
Impact of calreticulin overexpression on cells depleted of calreticulin. **(A)** Two-dimensional contour plots of BiP::sfGFP and mTurquoise signals (transfection marker) in parental cells and two derivative *CRT*Δ clones transiently transfected for 48 h and 72 h with plasmids encoding wild-type human CRT (CRT^WT^) or two lectin CRT mutants, CRT^Y92A^ and CRT^Y92A,W244A^. As a control for transfection, cells were transfected with a mock plasmid. Shadowed rectangles mark cells that were not selected for analysis because they had low levels of mTurquoise-tagged plasmid (indicating low transfection) or have very high transfection levels that could be toxic. Cells within unshadowed rectangles and marked with red delineate those cells expressing moderate/high levels of mTurquoise-tagged plasmid selected for analysis of BiP::sfGFP and XBP1s::mCherry signals distribution. Right histograms indicate the transfected, gated cells 72 h post-transfection (red rectangles). **(B)** Bar graph displaying the percentage of BiP::sfGFP^low^ cells (rescue phenotype) in mTurquoise-positive cells gated by the unshadowed boxes in “6A” as mean ± SD of data obtained from one (72 h) or two (48 h) independent experiments. Data from both *CRT*Δ clones have been combined in each time point of analysis. Statistical analysis was performed by a two-sided unpaired Welch’s t test and significance is indicated by asterisks (*p <0.05; *** p < 0.001; **** p < 0.0001; ns: non-significant). **(C)** CHO-K1 cells were transiently transfected with the hCRT plasmids used in “6A” for 72 h. Following transfection, cell lysates were recovered, and equal protein amounts were loaded into 12.5% SDS-PAGE gels and immunoblotted for hCRT to analyse protein stability in CRT mutants. GAPDH served as a loading control. The immunoblots are representative for two independent experiments.

### Calreticulin depletion exposes a negative feedback loop by which ATF6α represses IRE1

Given the well-established interplay between the ATF6α and IRE1 branches of the UPR, we sought to assess whether the constitutive activation of ATF6α observed in *CRT*Δ clones could be a contributing factor to the conspicuous downregulation of IRE1 activity. Through RT-PCR and gel electrophoresis, IRE1-mediated XBP1 mRNA splicing in parental cells and *CRT*Δ clones were compared. Under basal conditions, unspliced mRNA levels (XBP1u) were significantly higher in *CRT*Δ clones (Fig 7A left bar graph), consistent with the role of NATF6α in the transcriptional activation of XBP1 (Yoshida et al., 2001). In addition, compared to parental cells, *CRT*Δ cells exhibited a smaller fraction of spliced XBP1 mRNA (XBP1s) (Fig 7A right bar graph), consistent with the lower IRE1 activity measured by a XBP1s::mCherry reporter.

**Fig 7.**
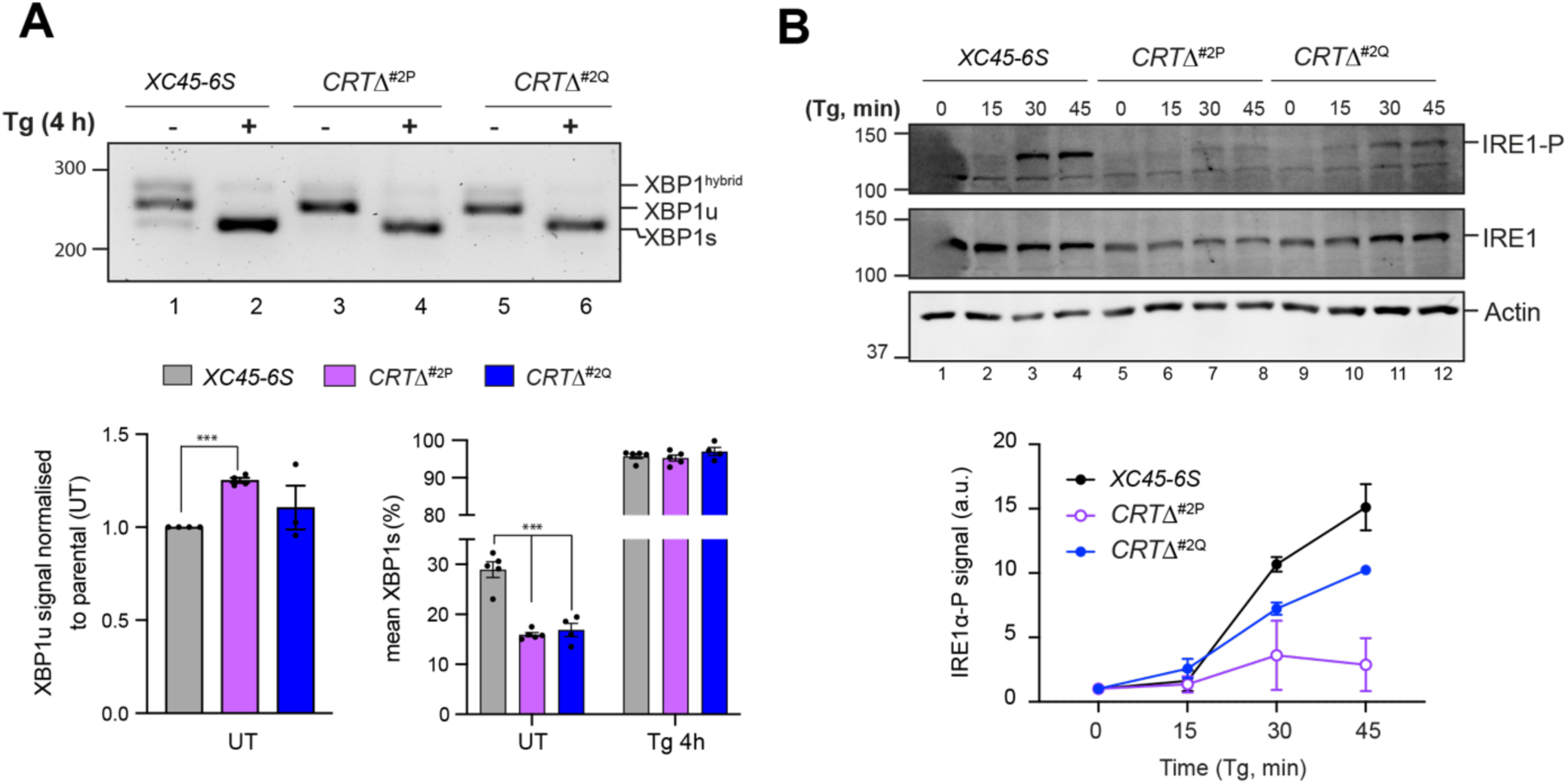
Calreticulin depletion exposes a negative feedback loop of ATF6α activity on IRE1. **(A)** *<u>Top panel.</u>* Agarose gel of XBP1 cDNA from thapsigargin (Tg)-treated parental cells and two *CRT*Δ clones. Migration of the unspliced (XBP1u), spliced XBP1 (XBP1s) and hybrid (XBP1^hybrid^) stained DNA fragments is indicated. The XBP1^hybrid^ band represents DNA species containing one strand of XBP1u and one strand of XBPs (Shang and Lehrman, 2004). Splicing was assessed by RT-PCR. *<u>Lower panel</u>*. Plots displaying the total XBP1u signal in the left panel, and the fraction of XBP1s in the right panel. For quantification, 50% of the hybrid band signal was assumed to be XBP1s and the other 50% XBP1u. The bars and error bars represent the mean ± SEM of data obtained from more than three independent experiments. Statistical analysis was performed by a two-sided unpaired Welch’s t test and significance is indicated by asterisks (*** p < 0.001). **(B)** Representative immunoblots of endogenous levels of cgIRE1 and phosphorylated IRE1 (IRE1-P) in parental cells and two *CRT*Δ clones basally and upon ER stress induction with 0.5 μM Tg for the indicated times. The levels of IRE1-P signal over the time in each genotype are represented in a graph as mean ± SD of data obtained from two independent experiments.

IRE1 activity is closely tied to its phosphorylation state (Shamu and Walter, 1996), therefore IRE1α-phosphorylation (IRE1α-P) levels were compared in parental and *CRT*Δ clones. Accordingly, *CRT*Δ cells exhibited lower levels of stress-induced phosphorylation of their endogenous IRE1 protein and slightly reduced total IRE1 levels (Fig 7B). These results supported earlier findings that IRE1 signalling is repressed by ATF6α activity (Walter et al., 2018). However, the defect in IRE1 activity in CRT-depleted cells was partial, as in the presence of ER stress, all genotypes exhibited a comparable increase in XBP1s (Fig 7A).

## Discussion

This study reports on a comprehensive and unbiased genome-wide CRISPR/Cas9-based knockout screen dedicated to discovering regulators of ATF6α. Whereas previous high-throughput studies have broadly studied modulators of the UPR (Adamson et al., 2016; Jonikas et al., 2009; Panganiban et al., 2019) we designed an ATF6α/IRE1 dual UPR reporter cell line to selectively detect regulators of ATF6α. Although a connection between CRT and ATF6α was proposed previously based on an interaction between overexpressed proteins (Hong et al., 2004), our study utilises unbiased genetic tools and manipulation of endogenous proteins to establish CRT’s role as a selective repressor of endogenous ATF6α signalling, mapping its function in retaining ATF6α in the ER.

It is notable that, though the screen identified well-known regulators of ATF6α processing (including S1P, S2P and NF-Y, and thus validating the experimental approach), only a limited number of new genes that contribute to ATF6α signalling emerged from the screen. Despite the biochemical evidence for the role of COPII-coated vesicles and the oligomeric state of ATF6α in its activation, no cluster of genes involved in COPII vesicular transport or PDIs were found among the most enriched pathways. Nor were any counterparts to SCAP or INSIG, regulators of ER-to-Golgi trafficking of the related SREBP proteins (Yang et al., 2002), identified. This might reflect essentiality of the genes involved or redundancy amongst them. Cell type specificity may also be implicated in failure to identify components that are unimportant to ATF6α activation in CHO-K1 cells. For example, deletion of ERp18, a small PDI-like protein implicated in ATF6α activation in HEK cells (Oka et al., 2019), affected neither the ATF6α nor IRE1α UPR branches in CHO-K1 cells studied here (Supplemental S6E).

The screen implicated two components of the nuclear pore complex (*NUP50* and *SEH1L*) in ATF6α pathway activity. While it might be tempting to consider a role for these components in the nuclear translocation of the processed soluble ATF6α N-terminal domain, there is no reason to think it would be regulatory, as it operates downstream of ER stress-regulated trafficking and processing events.

In addition to the identified S1P and S2P proteases, the *screen for ATF6*α *activators* identified FURIN, a Golgi-localised protease belonging to the subtilisin-like proprotein convertase family (like S1P protease) responsible for proteolytic activation of a wide array of precursor proteins within the secretory pathway (Braun and Sauter, 2019). FURIN deletion synergised with S1P inhibitors in attenuating ATF6α signalling. Additionally, ATF6α_LD has a conserved region (<u>R</u>T<u>K</u>S<u>RR</u>) related to the FURIN cleavage motif (generally described as RX-R-X-[K/R]-R↓ (Nakayama, 1997) (Supplemental Fig S4E). It has been suggested that S1P’s role in activation is to reduce the size of ATF6α’s_LD, thereby promoting S2P cleavage (Shen et al., 2002; Ye et al., 2000). FURIN cleavage could contribute to such a process and account for partial redundancy between S1P inhibition and FURIN depletion observed here. Nonetheless, *in vitro* FURIN failed to cleave the ATF6α_LD. These findings collectively suggest that FURIN may function as an indirect enhancer of ATF6α signalling, perhaps by setting up conditions in a post-ER compartment that favours ATF6α cleavage and activation.

The *screen for ATF6*α *repressors* revealed a substantial enrichment of genes involved in glycoprotein metabolism. Among that cluster, the most notable hit, not previously identified in other high-throughput UPR screens, was CRT, a well-characterised soluble ER chaperone with a key role in protein quality control. Intriguingly, its ER-membrane counterpart, CNX, did not emerge in our screen, despite the inclusion of six sgRNA targeting CNX in the CRISPR library. This suggests a special role of CRT in the regulation of ATF6α, at least in CHO-K1 cells. CRT may not be the only repressor of ATF6α; this role may be shared by BiP, which has been widely suggested to suppress all the three branches of the UPR under basal conditions. Nevertheless, because of BiP’s pervasive effects on ER proteostasis and its role in direct repression of the IRE1 branch, its role in regulating ATF6 might have been missed by this screen.

Disruption of CRT in the *XC45-6S* dual ATF6α/IRE1 UPR reporter cell line selectively activated the ATF6α pathway reporter under basal conditions. This activation was accompanied by an increased baseline trafficking of ATF6α from the ER to the Golgi, enhanced processing of ATF6α to its active form and increased endogenous BiP protein levels. Loss of CRT also resulted in a conspicuous basal downregulation of the XBP1s::mCherry reporter that was associated with a reduced activity of IRE1 in *CRT*Δ cells. These findings align with previous evidence that N-ATF6α suppresses IRE1 signalling (Walter et al., 2018), and highlights the importance of ATF6α/IRE1 crosstalk.

CRT engages monoglycosylated glycoproteins in the ER *via* its glycan-binding lectin domain, retaining them in the CRT/CNX cycle (Hebert et al., 1996; Ou et al., 1993). A role for CRT’s interaction with ATF6α_LD glycans in repressing ATF6α is suggested by the markedly diminished ability of a lectin mutant CRT^Y92A^ to repress the ATF6α pathway compared to the wild-type. However, it is worth noting that CRT has been observed to bind proteins also in a glycan-independent manner (Wijeyesakere et al., 2013). Both BLI data and immunoprecipitation from cell extracts revealed that CRT can associate with both wild-type and unglycosylated ATF6α (ΔGly). In the former, direct interaction is characterised by slow kinetics, consistent with previous measurements of glycosylation-independent binding of CRT to its clients (Table 1 from Wijeyesakere et al., 2013). These biophysical features, coupled with the residual ability of the lectin mutant CRT^Y92A^ to repress endogenous ATF6α, lead us to suggest that CRT repression of ATF6α may involve both glycosylation-dependent and independent interactions.

Currently, the structural basis for ATF6α_LD interactions with CRT remains unknown, and its role in the direct or indirect retention of ATF6α in the ER is a subject for speculation. Nonetheless, the observations presented here support previous studies indicating that the stable interaction of CRT with cellular substrates combines both lectin-dependent and lectin independent interactions into a hybrid mode. Moreover, our observations raise the possibility that the lectin activity of CRT is dispensable for its interaction with ATF6α_LD but implicated in recruitment of a third component required for repression. Either way titration of CRT by competing ligands is a plausible regulatory mechanism coupling ATF6α activity to the burden of unfolded (glycol) proteins in the ER (Fig 8).

**Fig 8.**
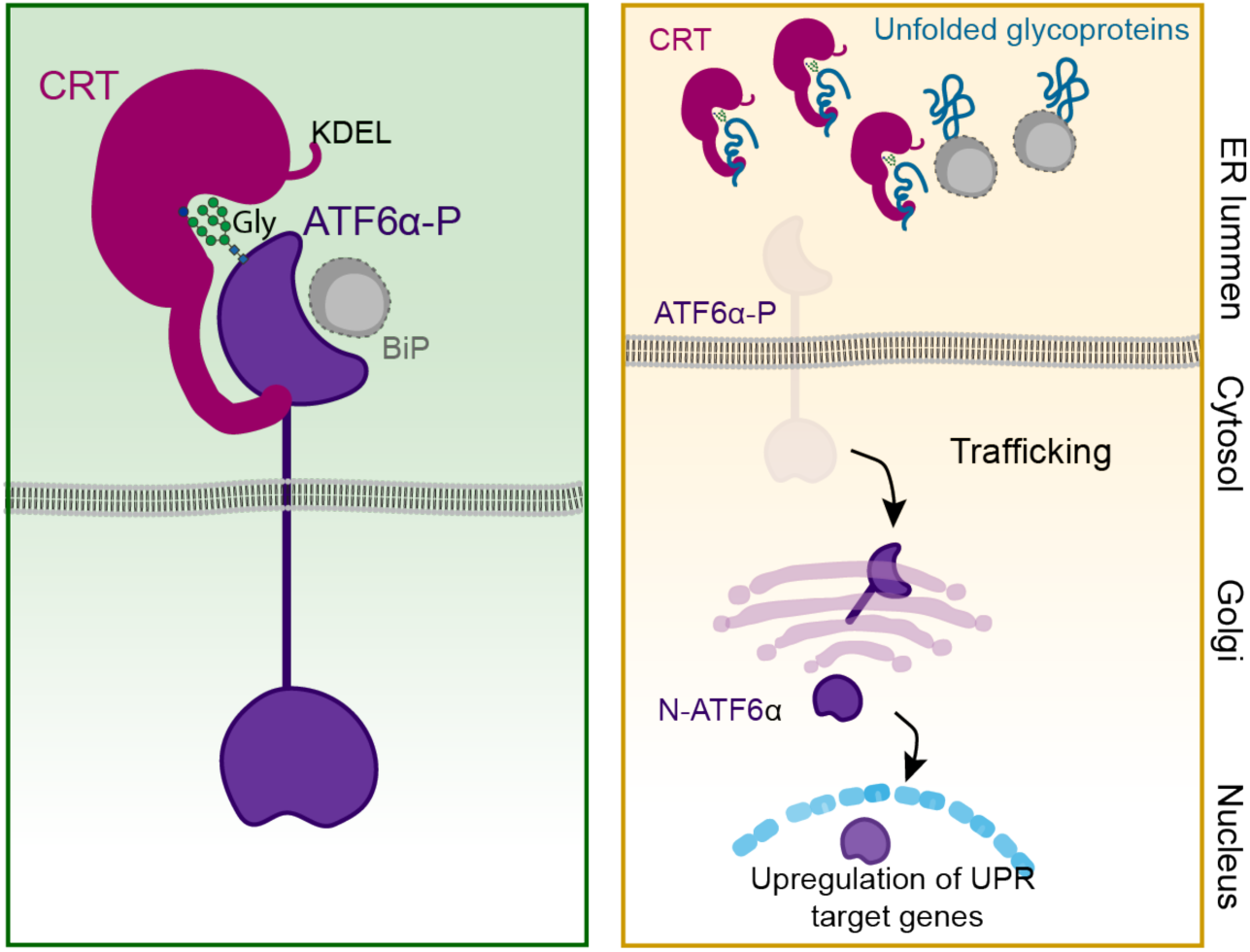
Proposed regulatory model of ATF6α by calreticulin (CRT) proteostasis. *<u>Left panel:</u>* When concentrations of competing ligands (unfolded glycoproteins) are low, ER retention of the ATF6α-CRT complex is favoured (*via* CRT’s KDEL signal). The ATF6α-CRT interaction is cartooned here as having both glycan (Gly)-dependent and glycan-independent components. *<u>Right panel</u>*: Unfolded glycoproteins compete with ATF6α for CRT and their increasing concentration favours ATF6α trafficking to the Golgi in stressed cells. A similar mechanism, involving BiP and shared by other UPR transducers, has been previously proposed (Bertolotti et al., 2000) and is included here as a complement to ATF6α-specific regulation by CRT. ATF6α-P refers to the precursor form of ATF6α, while N-ATF6α denotes the active form of ATF6α.

## Material and Methods

### Plasmids construction and reagents

Standard cloning techniques were used to create the recombinant DNA vectors listed in the Table S2. Single guide RNAs (sgRNAs) and oligonucleotides used in this study are listed in Table S3. Antibodies, reagents, and software are listed in Table S4.

### Mammalian cell culture, transfections, and treatments

Adherent Chinese Hamster Ovary (CHO-K1) cells (ATCC CCL-61) were maintained in Ham’s nutrient mixture F12 (Sigma) supplemented with 10% (v/v) serum (FetalClone-2, HyClone), 2 mM L-glutamine (Sigma), and 1% penicillin/streptomycin (Sigma). HEK293T cells (ATCC CRL-3216) were cultured in DMEM cell media (Sigma) supplemented as above. All cells were grown in a humidified 37°C incubator with 5% CO_2_ and passed every 2-3 days. When indicated, cells were treated with 5 μM Ceapin A7 (SML2330; Sigma), 15 μM S1P inhibitor (PF429242 dihydrochloride; Tocris), 2.5 μg/ml Tunicamycin (Tm, Melford), 4 mM 2-DeoxyD-glucose (2DG; ACROS Organics), 0.5 μM Thapsigargin (Tg; Calbiochem) and 16 μM 4μ8C (Sigma) for the indicated times. All compounds were diluted in pre-warmed culture medium and immediately added to the cells by medium exchange. Untreated cells were treated with DMSO solvent vehicle control. Transfections in CHO-K1 cells were performed using Lipofectamine LTX (Thermo Fisher Scientific, USA) at 1:3 DNA (µg) to LTX (µl) ratio, as for HEK293T using TransIT-293 Transfection Reagent (MIR2704, Mirus), according to the manufacturer’s instructions. Typically, cells were seeded at a density of 2.5 x 10^5^ cells/well in 6-well plates a day before transfection and analysed 48 - 72 h after transfection.

### Mammalian cell lysates and immunoblotting

CHO-K1 or HEK293T cells were cultured in 6-, 12-well plates or 10-cm dishes until reaching 95% confluence. Cells were washed twice with prechilled PBS on ice, and whole-cell extracts were scrapped out in 1mM EDTA-PBS, pelleted at 370*xg* for 10 min at 4°C and incubated in Nonidet-lysis buffer (150 mM NaCl, 50 mM Tris-HCl pH 7.5, 1% (v/v) NP-40) supplemented with 2x protease inhibitor cocktail (Roche Applied Science) for 30 min. Next, the samples were clarified at 21,130*xg* for 20 min at 4°C and the supernatants were transferred to fresh tubes. Protein concentration was determined using Bio-Rad protein assay. To assess the interaction between CRT and ATF6α through either Bio-Layer Interferometry (BLI) or co-immunoprecipitation, the lysis buffer was supplemented with 5mM CaCl_2_. The protein samples were separated on 8-12.5% SDS-PAGE under reducing conditions and transferred onto PVDF membranes as described previously (Ordóñez et al., 2013). Membranes were probed with the following primary antibodies: rabbit polyclonal anti-calreticulin (1:1000) (ab19261, abcam; to detect human CRT); chicken polyclonal anti-calreticulin (1:1000) (PA1-902A, Invitrogen; to detect Chinese hamster CRT); rabbit polyclonal anti-calnexin (1:1000) (SPA-860, Enzo Life Sciences); rabbit polyclonal anti-FURIN (1:1000) (ab3467, abcam); mouse monoclonal anti IRE1_LD serum (NY200) (1:1000) (Bertolotti et al., 2000); rabbit monoclonal anti p-IRE1 (1:1000) (Genetech); rabbit polyclonal anti GFP (NY1066) (1:1000) (Marciniak et al., 2004) made against purified bacterially expressed GFP following removal of the GST affinity tag *via* thrombin cleavage site; chicken anti-BiP IgY (1:1000) (Avezov et al., 2013); mouse monoclonal anti-FLAG M2 (1:1000) (F1804, Sigma); mouse monoclonal anti-KDEL (10C3) (1:1000) (ENZ-ABS679; Enzo Life Sciences); mouse monoclonal anti-actin (1:1000) (clone C4-691002, Fisher Scientific); rabbit monoclonal anti-GAPDH (1:2000) (14C10; Cell Signalling Technology); rabbit polyclonal anti-α/β-Tubulin (1:1000) (CST2148S, Cell Signalling Technology). For secondary antibody, IRDye fluorescently labelled antibodies or horseradish peroxidase (HRP) antibodies were used. Membranes were scanned using an Odyssey near infrared imager (LI-COR) and signals were quantified using ImageJ software. Signal quantification and analysis were performed using Prism 10 (GraphPad).

### Flow Cytometry and fluorescence-activated cell sorting (FACS)

To analyse UPR reporter activities, cells were grown on 6- or 12-well plates until reaching 8090 % confluence and then treated as indicated. For flow cytometry analysis, cells were washed twice in PBS, collected in PBS supplemented with 4mM EDTA, and 20,000 cells/sample were analysed by multi-channel flow cytometry on a LSRFortessa cell analyser (BD Biosciences). For FACS, cells were collected in PBS containing 4mM EDTA and 0.5% BSA and then sorted on a Beckman Coulter MoFlo cell sorter. Sorted cells were either collected in fresh media as a bulk of cells or individually sorted into 96-well plates and then expanded. Gating for live cells was based on FSC-A/SSC-A and for singlets was based on FSC-W/SSC-A. BiP::sfGFP and mGreenLantern (mGL) fluorescent signals were detected with an excitation laser at 488 nm and a 530/30 nm emission filter; XBP1s::mCherry fluorescence with an excitation laser 561 nm and a 610/20 nm emission filter, while mTurquoise and BFP fluorescence with an excitation laser 405 nm and a 450/50 nm filter emission. Data analysis was performed using FlowJo V10, and median reporter analysis was conducted using Prism 10 (GraphPad).

### Lentiviral production

Lentiviral particles were produced by co-transfecting HEK293T cells with the library plasmid (UK2561), the packaging plasmids psPAX2 (UK1701) and pMD2.G (UK1700) at 10:7.5:5 ratio using TransIT-293 Transfection Reagent (MIR2704, Mirus) according to the manufacturer’s instructions. Eighteen hours after transfection, medium was changed to medium supplemented with 1% BSA (Sigma). The supernatant containing the viral particles was collected 48 h after transfection, filtering through a 0.45 μm filter, and directly used to infect CHO-K1 cells seeded in 6-well plates for viral titration to calculate the amount of virus to use aiming a low multiplicity of infection (MOI) around 0.3 to ensure a single integration even per cell.

### Generation of the *XC45-6S* double UPR reporter cell by genome editing in *ROSA26* locus of CHO-K1 cells

The putative *ROSA26* locus in CHO cells has been recently identified and described as a “safeharbour” site for heterologous gene expression and stable long-term expression of a transgene (Gaidukov et al., 2018). To generate a double UPR reporter cell line, we used a previously-described *XC45* CHO-K1 cell line bearing a XBP1s::mCherry reporter expressing a pCAX-F-XBP1ΔDBD-mCherry transgene randomly integrated in the genome (Harding et al., 2019), for targeting the *ROSA26* locus for knock-in with a landing pad cassette, comprised the *cg*BiP promoter region containing ERSE-I elements, fused to the superfolded (sf) GFP into the CHO genome by CRISPR/Cas9 integration to make this reporter cell line also sensitive to ATF6α activity. The landing pad donor vector was constructed by appending homology arm sequences of *ROSA26* locus (440 bp predefined sequence from Gaidukov’s paper) to the 5’ promoter region of *cg*BiP (1,000 bp), along with the first 8 amino acids of the BiP CDS fused to the sfGFP. Homology arms were PCR amplified from CHO genomic DNA using predefined primers from Gaidukov’s paper. Subsequently, the *cg*BiP promoter, a geneblock containing sfGFP, and the flanking homology arms were fused by Gibson assembly cloning method, which allows multiple DNA fragments joining in a single, isothermal reaction (1 h at 50°C). Targeted integration was performed by co-transfection with the circular landing pad donor vector (UK2846) and a single sgRNA/Cas9 targeting *cgROSA26* locus (UK2847). Transfected cells were selected with puromycin (8 μg/ml) for 3 days and then treated with 2-Deoxy-D-glucose (4 mM) for 20 h to induce ER stress prior FACS. Cells were phenotypically sorted into 96-well plates by their activation of both UPR reporters (mCherry^high^; GFP ^high^). Phenotypically selected clones were subsequently confirmed by correct genetic integration. For stability integration, clones were maintained and expanded every 2-3 days for 1 month and weekly analysed with flow cytometry for mCherry and GFP expression under treatment with different ER stressors agents. For the wide-genome CRISPR/Cas9 screen, Cas9 was ultimately introduced by transduction with lentiviral particles prepared from packaging Lenti-Cas9 plasmid (UK1674). Cas9 activity was confirmed by targeting the exon 18 of IRE1 (*ERN1* locus) (UK2929) followed by induction of ER stress with 2-Deoxy-D-glucose (4 mM) for 20 h (Supplemental S1F).

### CRISPR/Cas9 screen

The screen was performed following established protocols using a pooled CHO-K1 knockout CRISPR library containing 125,030 sgRNAs targeting 21,896 genes, with six guides per gene, as well as 1,000 non-targeting sgRNAs as a negative control cloned into the lentiviral sgRNA expression vector pKLV-U6gRNA(BbsI)-PGKpuro2ABFP (Shalem et al., 2014; Ordóñez et al., 2021). Approximately 2.1 × 10^8^ CHO-K1 Cas9 stable cells carrying the dual UPR reporter ATF6α/IRE1 (*XC45-6S* cells) were infected at a multiplicity of infection (MOI) of 0.3 to favour infection with a single viral particle per cell. Two days after infection, transduced cells were selected with 8 µg/ml puromycin for 14 days. Afterwards, transduced cells were split into two subpopulations: one was treated with 2-Deoxy-D-glucose to induce ER stress, while the other one remained untreated. For the *ATF6*α *activator screen,* conducted under ER stress conditions, less than 2% of cells that showed a XBP1s::mCherry^high^; BiP::sfGFP^low^ phenotype were selected for analysis. Similarly, for the *ATF6*α *repressor screen,* conducted under basal (untreated) conditions, less than 2% of total sorted cells that showed a XBP1s::mCherry^low^; BiP::sfGFP^high^ phenotype were selected for analysis. Rounds of enrichment were carried out through cellular recovery, expansion, and sorting. Equal number of transduced cells, both untreated or 2-Deoxy-D-glucose treated, were passed without sorting as a control group in each round of sorting. To prepare samples for deep sequencing, genomic DNA from enriched and sorted populations as well as unsorted cells (to represent the entire library) was extracted from ∼3.6× 10^7^ and ∼1-3× 10^6^ cells, respectively, by incubation in proteinase K solution [100 mM Tris-HCl pH 8.5, 5 mM EDTA, 200 mM NaCl, 0.25% (w/v) SDS, 0.2 mg/ml Proteinase K] overnight at 50°C. Integrated sgRNA sequences were amplified by nested PCR and the necessary adaptors for Illumina sequencing were introduced at the final round of amplification. The purified products were quantified and sequenced on an Illumina NovaSeq 6000 by 50-bp single-end sequencing. Downstream analyses to obtain sgRNA read counts, gene rankings, and statistics were performed with MAGeCK software (Li et al., 2014). Gene ontology analyses were performed using Metascape with default parameters.

### Knockout cells using CRISPR/Cas9 technology

sgRNAs targeting exon regions of *Cricetulus griseus CRT*, *CNX*, *FURIN* and *ATF6*α were designed using CRISPR-Design tools from *CCTop* (https://cctop.cos.uniheidelberg.de:8043/index.html) and *CRISPy-Cas9 target finder for CHO-K1* (http://staff.biosustain.dtu.dk/laeb/crispy/) databases. sgRNAs were then cloned into the pSpCas9(BB)-2A-mTurquoise (UK2915) or pSpCas9(BB)-2A-Puro plasmids (UK1367) as previously reported (Ran et al., 2013). The integrity of the constructs was confirmed by DNA sequencing. To generate the knockout cells, 0.75-1 µg of sgRNA/Cas9 plasmids was transfected into *XC45-6S* cells using Lipofectamine LTX (Thermofisher) in a 6-well plate format. After 48 h post-transfection, mTurquoise^high^ cells were sorted into 96-well plates at 1 cell/well using a MoFlo Cell Sorter (Beckman Coulter). For cells transfected with sgRNA/Puro plasmids, selection was performed in the presence of 8 μg/ml puromycin, and serial dilution was used to isolate single clones. The knockouts cells were confirmed by Sanger sequencing and immunoblotting.

### ATF6α tagged cells by CRISPR/Cas9 genome-editing

To investigate potential modulators of ATF6α activity following the CRISPR/Cas9 high-throughput screen using biochemistry and microscopy-compatible tools, two mammalian cell lines were engineered. These cell lines either harboured a knock-in of the endogenous *cgATF6*α locus or carried a stable transgene encoding ATF6α_LD tagged with a bright GFP. The tags were positioned at the N-cytosolic terminus, known to be well tolerated.

### 3xFLAG-mGreenLantern_ATF6α knock-in cells (*2K* cells)

Protein alignment of *hs*ATF6α and *cg*ATF6α (UniProt database) indicated that translation initiation predominantly occurs at the second Met at the beginning of exon 2 (<u>M</u>ESP). Instead of tagging at the first Met in the CHO genome, tags were introduced before the third Met. CHO-K1 plain cells were offered with a unique pSpCas9(BB)-2A-mCherry targeting exon 2 of *cg*ATF6α locus (UK2942) and a donor plasmid containing homology arms flanking a minigene encoding a 3xFLAGmGreenLantern(mGL)-TEV sequence geneblock (UK2955) to create an endogenous tagged ATF6α. After 48 h post-transfection, mGL^high^ cells were sorted as single cells into 96-well plates. Clones that preserved stable mGL fluorescent signal by flow cytometry and showed expected ATF6α processing upon ER stress induction (DTT) were subsequently confirmed by correct genetic integration. Microscopy characterisation was performed under basal conditions or after treatments with ER stressors (DTT), Ceapin A7 and S1P inhibitor. A clone referred to as *2K* was expanded and selected for functional analysis as a 3xFLAG-mGL tagged endogenous ATF6α.

### GFP-cgATF6α_LD^S1P,S2Pmut^ (*UK3122* cells)

ATF6 has a half-life of about 3 h (George et al., 2020). To increase the proportion of intact, active N-ATF6α molecules reaching the Golgi, a genetic construct featured the TM and LD of ATF6α, deliberately lacking S1P and S2P cleavage sites, and further strategically appended with an Avi-tag at the C-terminal, was engineered. This construct was then used for a dual purpose: as a tool for both microscopy studies as well as to be biotinylated and purified to be used as a ligand in BLI experiments. Shen and Ye’s research provided the basis for the design of S2P and S1P mutations (N391F & P394L for S2P mutant, and LL408/9VV for S1P mutant) (Shen and Prywes, 2004; Ye et al., 2000). A DNA-plasmid encoding eGFP_cgATF6α_TEV_LD410-659_S12PMUT_AviTag_pCEFL_pu (UK3122) was introduced into CHO-K1 plain cells *via* Lipofectamine LTX. Transfected cells were selected for resistance to puromycin (8 µg/ml), and the isogenic clones were selected for confocal microscopy assays and *in vitro* BLI interaction assays.

### A genetic platform to generate cgATF6**α_**LD variants knock-in cells

#### ATF6αΔ

Two sgRNA sequences targeting exon 9 and 10 of cg*ATF6*α were incorporated into pSpCas9(BB)-2A-mTurquoise plasmid (UK2915) to generate two sgRNA/Cas9 expression plasmids (UK3021 and UK3024) according to the published procedure (Ran et al., 2013). These plasmids were then transfected into CHO-K1 *XC45-6S* double UPR reporter cells. After 72 h, mTurquoise^high^ cells were sorted as single cells into 96-well plates. Genomic DNA from clones showing no response to ATF6α activation (BiP::sfGFP^low^) upon tunicamycin treatment (0.5 µM, 20h) was extracted for Sanger sequencing. Clones with frame shifts causing deletions in both alleles of the DNA region flanking by both sgRNAs were selected for further functional assays.

#### cgATF6α_LD “minigenes” knock-in

A unique sgRNA sequence targeting the exon 10 of α*ATF6*α**Δ** cells was inserted into the pSpCas9(BB)-2A-mTurquoise plasmid to create the sgRNA/Cas9 plasmid UK3025. Three repair “minigenes” templates were constructed: A wildtype, glycan free (ΔGly) or a cysteine free (ΔC) LD to be reconstituted in ATF6αΔ cells by CRISPR/Cas9-mediated homology-directed repair (HDR).

For the construction of the wild-type LD repair template (WT; UK3054), a minigene block that contains exons 10 −17 of *cgATF6*α (785bp; GenScript) was digested with *Age*I and *Afl*II and ligated into *Age*I/ *Afl*II-digested pUC57 plasmid. To create a Glycan free LD (ΔGly; UK3086) template, a minigene block containing exon 10 −17 (GenScrip) with the three N-glycosylation sites were mutated by replacing Thr 464, 575 and 634 with Ala, was cloned into pUC57 plasmid as previously described. Similarly, to generate a Cysteine-free LD (ΔC; UK3147) template, a minigene block containing exon 10 −17 (GenScrip) with Cys457 and Cys607 replaced with Ser, was cloned into pUC57 plasmid as previously performed. The integrity of the constructs was confirmed by DNA sequencing.

### Immunoprecipitation using GFP-Trap Agarose

Equal volumes of the cleared and normalised lysates were incubated with 15-20 μl GFP-Trap Agarose (ChromoTek) pre-equilibrated in lysis buffer. The mixture was incubated overnight at 4 °C with rotation. The beads were then recovered by centrifugation (845*xg*, 3 min) and washed four times with washing buffer (50 mM Tris-HCl, 1 mM EDTA, 0.1% Triton X-100) and one more with PBS. Proteins were eluted in 35 μl of 2×SDS-DTT sample for 5 min at 95°C. Equal sample volumes were analysed by SDS-PAGE and immunoblotting, as described above. Normalised cell lysates (15-30 μg) were loaded as an ‘input’ control.

### Bio-layer Interferometry (BLI), protein expression and *in vivo* biotinylation

BLI studies were undertaken using on the Octet RED96 System (Pall FortéBio) using ATF6α_LD variants as biotinylated ligands that were immobilised on streptavidin (SA)-coated biosensors (Pall FortéBio) and rat calreticulin (*r*CRT) as the analyte. Prior to use, SA-coated biosensors were hydrated in a BLI buffer (150 mM NaCl, 20 mM Tris, 1mM CaCl_2_) for 10 min. BLI reactions were prepared in 200 μl volumes in 96-well microplates (greiner bio-one), at 30°C. Ligand loading was performed for 600 sec until a binding signal of 1 nm displacement was reached. Following ligand immobilisation, the sensor was baselined in the reaction buffer for at least 200 seconds, after which it was immersed in wells containing the analyte for association for 1200 seconds. After each association, the sensor was dipped into a buffer containing well to allow dissociation for 2400 sec.

Biotinylated ATF6αLD constructs [ATF6αLD^WT^ (UK3122); ATF6αLD^ΔGly^ (UK3141) and ATF6LD^ΔC^ (UK3157)] were expressed in HEK293T cells, each one encoding a N-terminal GFP tag followed by a TEV protease cleavage site just after the TM and an Avi-tag at Cterminal. S1P and S2P cleavage sites were also mutated to increase expression yield. Biotinylation was conducted *in vivo*, where cells were co-transfected with a BirA-encoding plasmid (biotin ligase, UK2969) in a 1:0.06 ratio. After 24 h, media was replaced with fresh media containing 50 μM biotin dissolved in DMSO and cells were harvested and lysed the following day. The lysis buffer was supplemented with 5 mM CaCl_2_. Post immunoprecipitation using GFP-Trap Agarose, proteins were eluted by incubation with an excess of TEV protease (UK759, 16 μg) in 400 ul TEV cleavage buffer (25 mM Tris–HCl pH 7.4, 150 mM NaCl, 0.05% NP40, 0.05mM CaCl_2_, 0.05 mM TCEP) for 16 h at 4°C with orbital rotation. Cleavage products were snap-frozen in liquid nitrogen and stored at −80°C for BLI experiments.

Wild-type full-length *r*CRT (UK3142) was expressed in *E.Coli* cells (BL21 C3013). Bacterial cultures were grown in LB medium with 100 µg/ml ampicillin at 37°C to an OD_600nm_ of 0.60.8. Expression was induced with 1 mM IPTG, and the cells were further grown for 16 h at 18°C. After cell sedimentation by centrifugation, the pellets were lysed with a high-pressure homogeniser (EmulsiFlex-C3; Avestin) in bacterial lysis buffer (50 mM Tris-HCl pH 7.4, 500 mM NaCl, 2mM CaCl_2_, 20mM imidzole) containing protease inhibitors (1 mM PMSF, 2 µg/ml pepstatin, 8 µg/ml aprotinin, 2 µM leupeptin) and 0.1 mg/ml DNaseI. Typical yields from 2 litres of bacterial culture were obtained.

### Immunoprecipitation using GFP-Trap Agarose

To investigate the interaction between CRT and ATF6α in cells, HEK293T cells were cultured in 10-cm dishes (3-4 plates per sample) and transfected with constructs expressing either GFPATF6α_LD^WT^ (UK3122) or GFP-ATF6α_LD^ΔGly^ (UK3141). Equal volumes of the cleared and normalised lysates were incubated with 15-20 μl GFP-Trap Agarose (ChromoTek) preequilibrated in lysis buffer. The mixture was incubated overnight at 4°C with rotation. The beads were then recovered by centrifugation (845*xg*, 3 min) and washed four times with washing buffer (50 mM Tris-HCl, 1 mM EDTA, 0.1% Triton X-100). Proteins were eluted in 35 μl of 2×SDS-DTT sample for 5 min at 95°C. Equal sample volumes were analysed by SDSPAGE and immunoblotting, as described above. Normalised cell lysates (15-30 μg) were loaded as an ‘input’ control.

### Analysis of endogenous ATF6α processing in CRT-depleted cells

To investigate the impact of CRT depletion on ATF6α cleavage, CRT was depleted in *2K* cells (3xFLAG-mGL-ATF6α) by CRISPR/Cas9 gene editing (UK2991). *2K-CRT*Δ derivative clones were confirmed by Sanger sequencing and selected for further experiments. Both *2K* parental cells and *2K-CRT*Δ derivative cells were cultured in 10-cm dishes for 2 days and then treated with 2-Deoxy-D-glucose (4 mM) for a time course up to 6 h before harvested. Following treatment, the cells were lysed in Nonidet-lysis buffer for extracting the cytosolic fraction, as previously described. For nuclear extraction, high-salt lysis buffer (500 mM NaCl, 50 mM Tris-HCl pH 7.5, 1% (v/v) NP-40) supplemented with 2x protease inhibitor cocktail was added to the nuclear pellets, followed by rotation at 4°C for 2 h. Subsequently, samples were centrifuged at 15,000 rpm at 4°C for 15 min, and the soluble portion containing the nuclear extract was combined with the cytosolic fraction. Equal volumes of clarified and normalised cytosolic and nuclear extracts were incubated with 15-20 μl GFP-Trap Agarose (ChromoTek) for overnight at 4 °C with rotation. Afterwards, beads were recovered by centrifugation, washed three times in washing buffer and eluted in 35 μl of 2×SDS-DTT sample for 5 min at 95°C.

### Live cell confocal microscopy

Cells were grown on 35 mm live cells imaging dishes (MatTek), transfected as required, and maintained in culture for 48 h after transfection. Live cell microscopy was conducted using an inverted confocal laser scanning microscope (Zeiss LSM880) in a 37°C/5% v/v chamber with a 64x oil immersion objective lens with 1 Airy unit pinhole using the 488 nm (GFP; mGL), 405 nm (DAPI) and/or 594 nm (mCherry) lasers as appropriate. Co-localisation analysis between fluorescence channels (Pearson correlation coefficient) within individual cells was performed using Volocity software, version 6.3 (PerkinElmer).

### RNA isolation and reverse transcription

To extract total RNA, cells were treated with TRIzol™ Reagent (Invitrogen) for 10 min and then transferred to a new tube. Chloroform (200 µl) was added into each sample, vortexed for 1 min, and then centrifuged at 13,500*xg*, at 4°C for 15 min. The upper clear phase was collected and mixed with equal volume of 70% ethanol. Afterward, PureLink™ RNA mini kit (Invitrogen) was used for washing and elution. RNA yield was assessed using NanoDrop, and 2 µg of RNA from each sample was evaluated for integrity *via* electrophoresis on a 1.2% agarose gel.

For reverse transcription (RT)-PCR, 2 µg of RNA was initially heated up at 70°C with RT buffer (Thermo Scientific; Cat #EP0441), 0.5 mM dNTP and 0.05 mM Oligo dT18 for 10 min. Subsequently, the reaction was supplemented with 0.5 µl RevertAid Reverse Transcriptase (Thermo Scientific) and 100 mM DTT, followed by incubation at 42°C for 90 min. The resulting cDNA product was diluted up to 1:4.

### PCR analysis of XBP1 mRNA splicing

XBP1s and XBP1u fragments were amplified from cDNA by PCR using primers flanking the splicing site identified by IRE1 with NEB Q5® High-Fidelity 2X Master Mix, as previously reported (Neidhardt et al., 2023). Briefly, XBP1u (255bp) and XBP1s (229bp) were separated in a 3 % agarose gel by electrophoresis and stained with SYBR Safe nucleic acid gel stain (Invitrogen). Additionally, a XBP1^hybrid^ band (280 bp), was observed. The percentage of XBP1s was quantified by determining the band intensity with Fiji, v1.53c. For quantification, it was assumed that 50% of the XBP1^hybrid^ signal represented XBP1s, while the remaining 50% represented XBP1u.

### Immunoprecipitation and FURIN cleavage assay

To investigate whether ATF6α is directly cleaved by FURIN, GFP-ATF6α_LD^S1P,S2Pmut^ (UK3122) construct was expressed and purified from a *FURIN*Δ clone (#2.2.G). Equal volumes of clarified and normalised cell lysates were incubated with 15-20 μl GFP-Trap Agarose (ChromoTek) for overnight at 4 °C with rotation. Afterwards, beads were recovered by centrifugation, washed three times in washing buffer and then incubated with 75 μl of FURIN cleavage buffer (20 mM HEPES, 1 mM CaCl_2_, 0.2 mM β-mercaptoethanol, 0.1% Triton X-100; pH 7.5) supplemented with 1 U FURIN (#P8077S, NEB) for 6 h at 25°C. As a negative control, calcium-depleted FURIN buffer was used. Additionally, a FURIN control substrate, MBP5-FN-paramyosin-ΔSal (#E8052S, NEB) containing a FURIN site, was included in parallel reactions and incubated for 6h at 25°C. The eluted protein was then analysed either *via* immunoblot or Coomassie blue staining.

### Analysis of ATF6α Redox Status

Both *2K-*parental cells and *2K-CRT*Δ derivative cells were cultured in 10-cm dishes for 2 days and then treated with 2-Deoxy-D-glucose (4 mM) for 3 h before harvested. Following treatment, cells were harvested in PBS containing 1mM EDTA and 20 mM NEM and lysed in Nonidet-lysis buffer supplemented with 20 mM NEM. All subsequent buffers were also supplemented with 20 mM NEM. Equal volumes of clarified and normalised cell lysates were incubated with 15-20 μl GFP-Trap Agarose (ChromoTek) for overnight at 4°C with rotation. Afterwards, beads were recovered by centrifugation, washed three times in washing buffer/NEM and eluted in SDS/NEM loading buffer. After heating at 95°C, eluted proteins were split into two equal samples and separated under reducing (+ 50 mM DTT) or non-reducing SDS/PAGE, followed by immunoblot.

### Statistics

Data groups were analysed as described in the figure legends using GraphPad Prism10 software. Differences between groups were considered statistically significant if * p < 0.05; ** p < 0.01; *** P < 0.001; and **** p < 0.0001. All error bars represent mean ± SEM.

## Data availability

The raw and processed high-throughput sequencing data reported in this paper have been uploaded in NCBI’s Gene Expression Omnibus (GEO, accession number: GSE254745).

## Acknowledgments

We thank the CIMR flow cytometry core facility team (Reiner Schulte, Chiara Cossetti and Gabriela Grondys-Kotarba), the microscopy team (Matthew Gratian and Mark Bowen) for technical support and the Huntington lab for access to the Octet machine. We also thank Marcella Ma (CRUK) for assistance with NGS and Avi Ashkenazi (Genentech) for the monoclonal antibody against human IRE1_LD. This work was supported by Wellcome Trust Principal Research Fellowship to DR (Wellcome 224407/Z/21Z),

## Author’s contributions

**AO:** co-led, conceptualisation; conceived and initiated the project; designed and conducted the CRISPR/Cas9 screen; conducted *in vivo* experiments and biophysical experiments; analysed and interpreted the data; prepared figures and wrote the original draft. **JT:** designed and conducted most of the *in vivo* experiments; analysed and interpreted the data; contributed to discussion. **LH:** conducted *in vivo* experiments; analysed and interpreted the data; **GG:** conducted *in vivo* experiments; interpreted the data; contributed to discussion. **HPH:** constructed the XCdel45 precursor cell line; interpreted the data; contributed to discussion. **DR:** conceptualisation; supervision; funding acquisition; investigation; interpreted data; cowrote the manuscript. All authors read and approved the final manuscript.

## Supplemental Figures

**Supplemental Fig S1.**
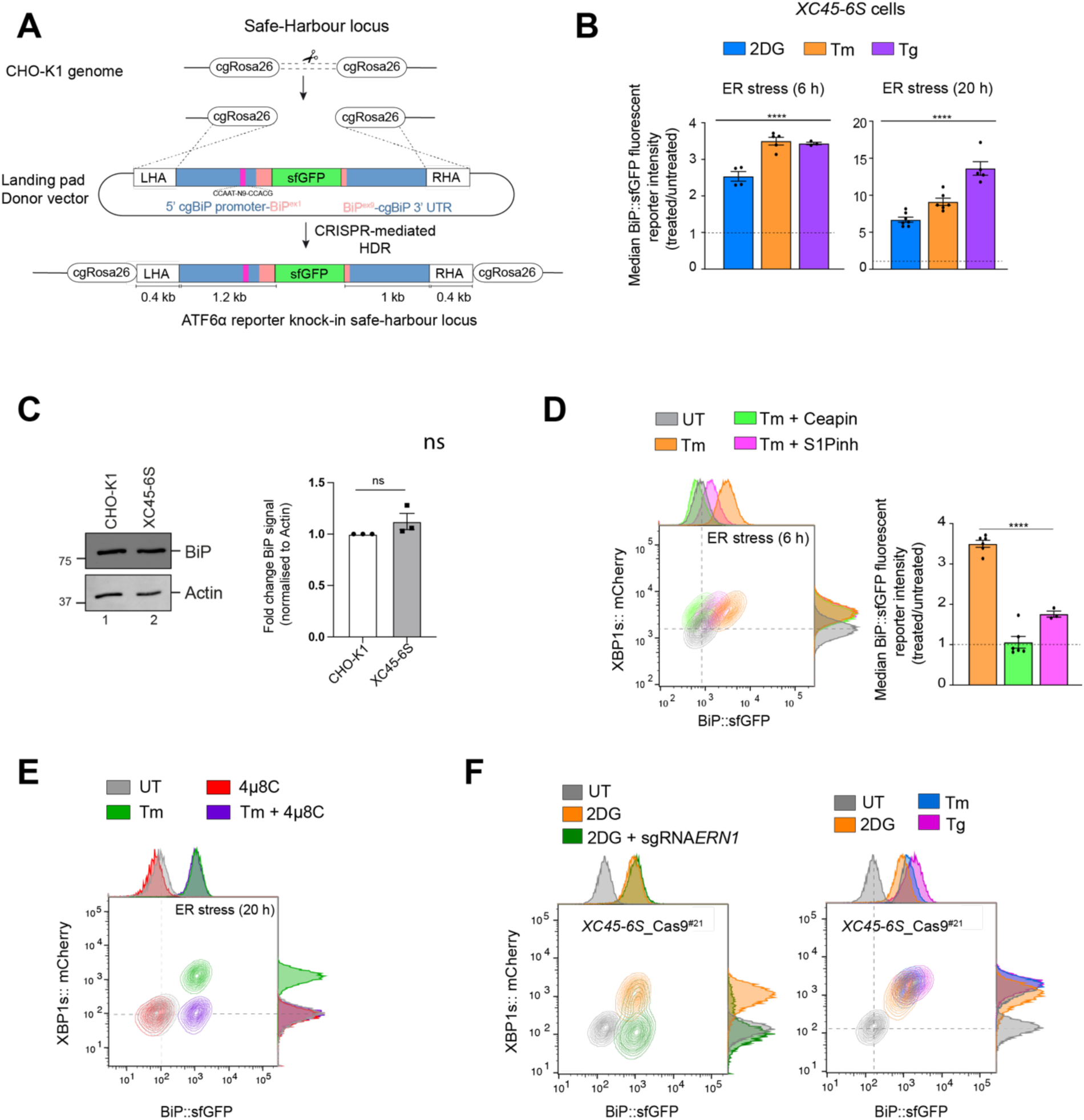
Characterisation of the ATF6α/IRE1 dual UPR reporter cell (*XC45-6S*). **(A)** Schematic representation of the strategy for targeted integration at the *cgROSA26* safe-harbour locus in the CHO genome, mediated by CRISPR/Cas9-mediated homology-directed repair (HDR). The approach involves an unique sgRNA/Cas9 sequence targeting the *cgROSA26* locus (Gaidukov et al., 2018) and a landing pad donor plasmid containing homology arms (LHA and RHA) flanking a minigene encoding the promoter region of the *cgBiP* fused with the superfolder green fluorescent protein (sfGFP) to create a BiP::sfGFP reporter to monitor ATF6α activity in cells. **(B)** <u>Related to</u> Fig 1A<u>, top panel</u>. Quantification of fold-change BiP::sfGFP signals in *XC45-6S* dual UPR reporter cells under basal conditions (untreated) and treated with the ER stressors 2DG, Tm and Tg (6 h or 20 h). Shown is the normalised (treated/untreated) median fluorescence intensity (MFI) as mean ± SEM of independent repetitions (n > 3). **(C)** Representative immunoblots of endogenous BiP protein levels in cell lysates from parental CHO-K1 cells and ATF6α/IRE1 dual UPR reporter cells (*XC45-6S*). The samples were also blotted for actin (loading control). The right graph bar shows the fold change of BiP signal normalised to the signal of the loading control in three independent experiments indicated by mean ± SEM. **(D)** Two-dimensional contour plots of BiP::sfGFP and XBP1s::mCherry levels in *XC45-6S* cells under basal conditions (UT, grey) or after treatment with tunicamycin alone (2.5 µg/ml, Tm, orange) or in combination with Ceapin A7 (5 µM, green) or S1P inhibitor (15 µM, pink) for 6 h. Histograms of the signal in each channel are displayed on the axes. The bar graph shows the mean ± SEM of the normalised median BiP::sfGFP reporter intensity from more than four independent experiments. **(E)** Contour plot, as in “S1D”, from parental cells treated with tunicamycin alone (Tm, 2.5 µg/ml, green) or in combination with the IRE1 inhibitor 4µ8C (10 µM, purple) for 18 h. A representative dataset from two independent experiments is shown. **(F)** Characterisation of the ATF6α/IRE1 dual UPR reporter cell line used in the CRISPR/Cas9 screen after stable integration of Cas9. *<u>Left panel.</u>* Contour plots to confirm Cas9 stable integration. Cells were analysed under basal conditions (grey) or after treatment with 2DG alone (orange) or transfection with an individual lentiGuide-Puro plasmid (without expression of Cas9) targeting exon 18 of IRE1 (encoded by *cgERN1*, green). ER stress treatments with 4 mM 2DG lasted 24 h. *<u>Right panel</u>*. As in “left panel”, but cells were treated with three different ER stressors for 20 h: 2DG (4 mM), Tm (2.5 µg/ml) or Tg (0.5 µM). A representative dataset from two independent experiments is shown. All statistical analysis was performed by a two-sided unpaired Welch’s t-test and significance is indicated by asterisks (**** p < 0.0001; ns: non-significant).

**Supplemental Fig S2.**
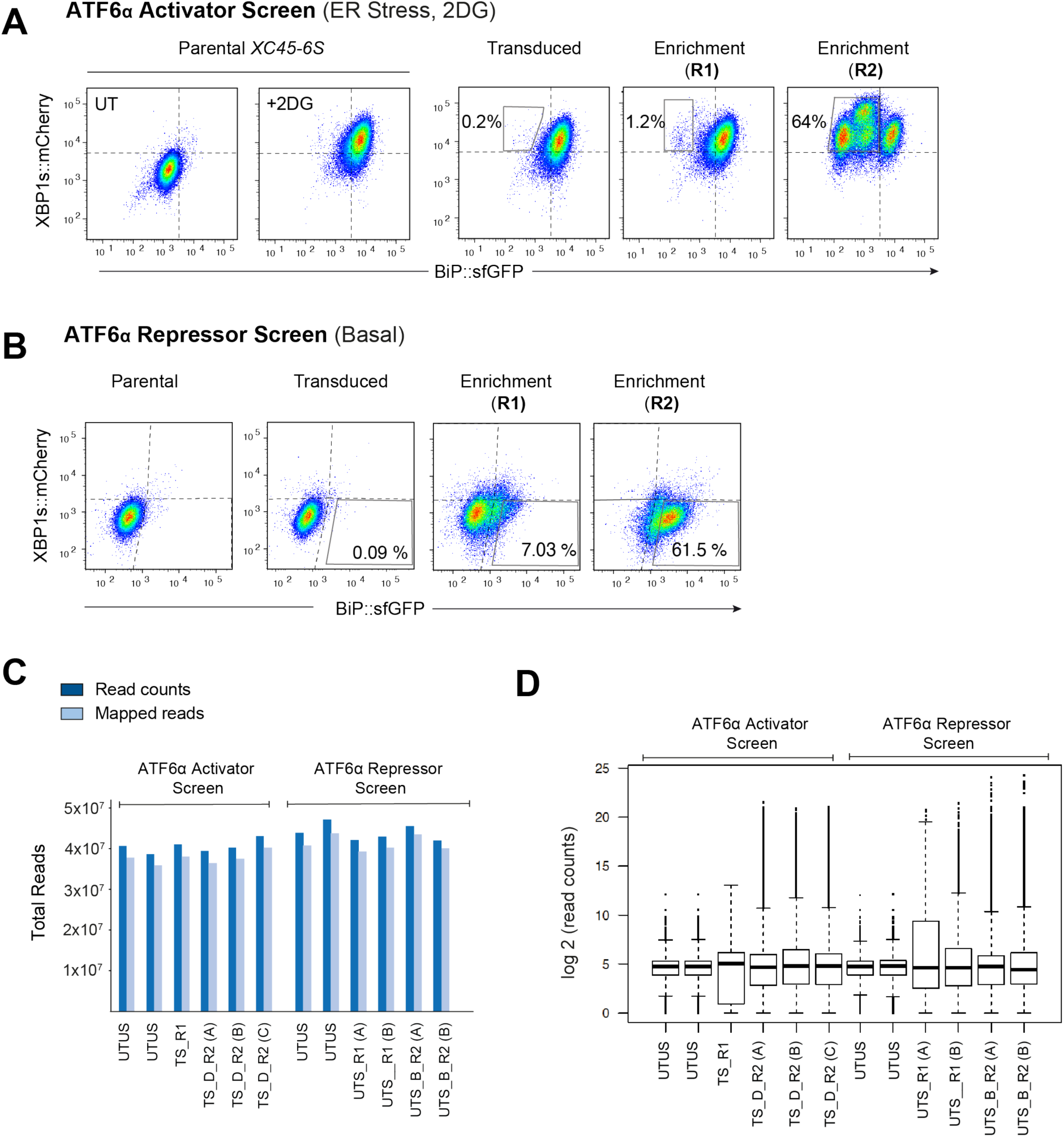
Enrichment method and quality control data analysis by MAGeCK of the CRISPR/Cas9 screens. **(A)** Two-dimensional contour plots of BiP::sfGFP and XBP1s::mCherry signals in *XC45-6S* obtained after each sort of enrichment (R1: round 1; R2: round 2) in the screen designed to identify ATF6α *activators*. The plots reveals a progressive enrichment of cells displaying CRISPR/Cas9-induced genetic perturbations that specifically impairing BiP::sfGFP activation upon UPR induction with 2DG (4 mM). **(B)** Contour plots, as in “S2A”, representing the screen designed to identify ATF6α *repressors* by successive rounds of enrichments. The focus is on cells exhibiting CRISPR/Cas9-induced genetic perturbations that lead to the constitutive activation of the BiP::sfGFP reporter under basal conditions, while leaving the XBP1s::mCherry reporter unaffected. **(C)** Analysis of total read counts and mapped reads to the CHO library analysed by MAGeCK [UTUS: untreated and unsorted; TS: treated (plus 2DG) and sorted; R1: Round 1 of enrichment; R2: Round 2 of enrichment; D: Dull GFP population; B: Bright GFP population; A-C: pools of cells].**(D)** Frequency distribution of sgRNA in each sample, showing the median-normalised read counts.

**Supplemental Fig S3.**
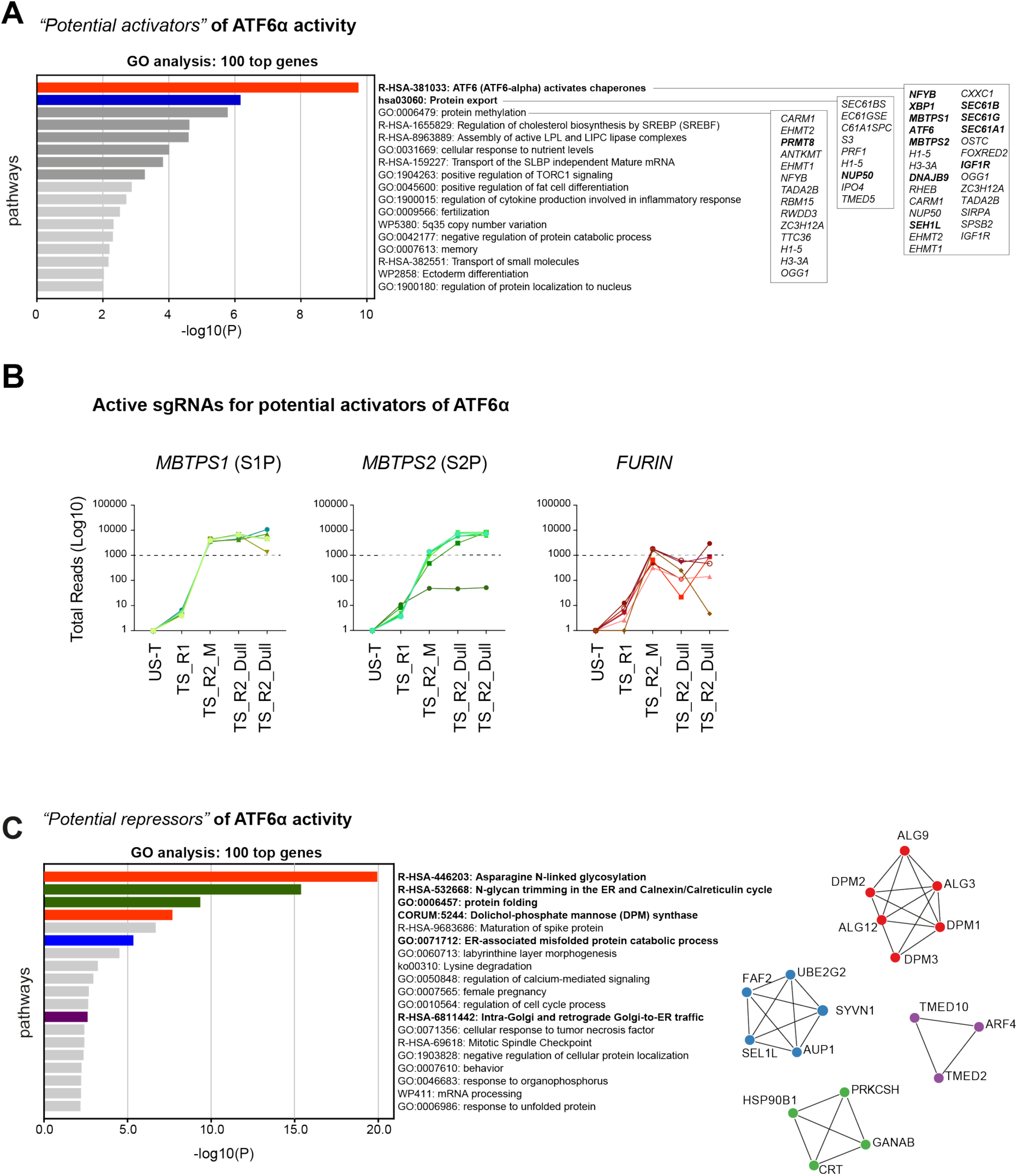
Integration of CRISPR screen hits. **(A)** Pathway enrichment analysis for biological processes using Gene Ontology (GO) terms for the top 100 hits identified in the CRISPR/Cas9 screen aimed to identify potential activators of ATF6α signalling. Selected pathways are ranked by p-value. **(B)** Total reads count enrichment through the selection process for each active sgRNA targeting the most significant genes identified in the *activator ATF6*α *screen*, including known regulators of ATF6α (*MBTPS1-*S1P and *MBPTPS2-*S2P) and those with possible roles (*FURIN*) in the activation of ATF6α signalling. **(C)** Pathway enrichment analysis, as in “S3A”, for the top 100 hits identified in the CRISPR/Cas9 screen aimed to identify potential *repressors* of ATF6α signalling. Selected pathways are ranked by p-value. On the left, a protein-protein interaction network (Metascape) for the 100 top hits identified in the CRISPR/Cas9 screen for potential repressors of ATF6α signalling is shown.

**Supplemental Fig S4.**
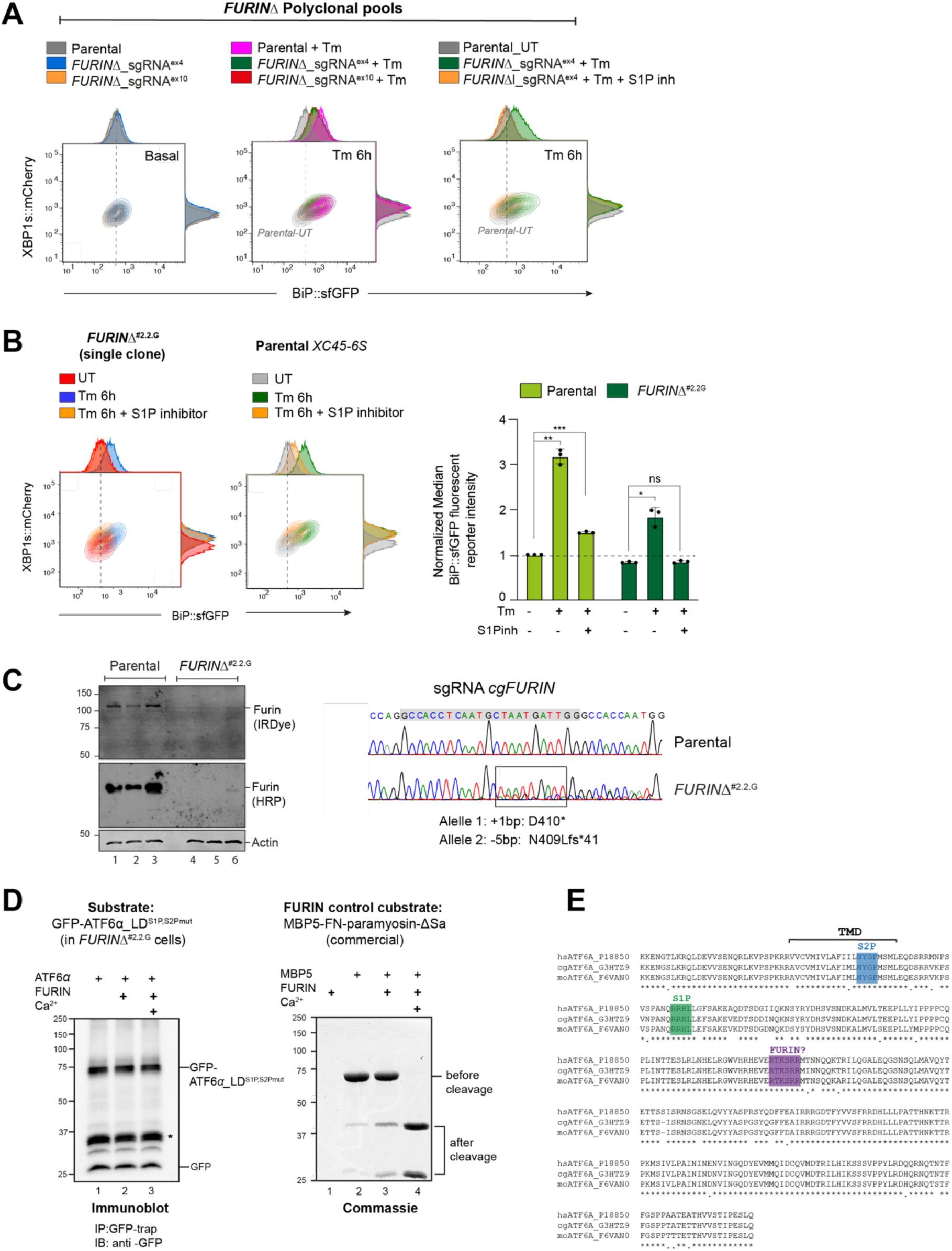
FURIN depletion decreases responsiveness to ATF6α activation upon ER stress. **(A)** Two-dimensional contour plots of BiP::sfGFP and XBP1s::mCherry signals in a sorted pooled population with FURIN depletion through CRISPR/Cas9 sgRNAs targeting exon 4 and 10. The analysis was conducted under basal conditions (left panel), after tunicamycin (Tm)-ER stress induction alone for 6h (middle panel) or in combination with S1P inhibitor treatment (15 µM) (right panel). Parental cells were used as a reference. A representative dataset from more than three experiments is shown. **(B)** Contour plots as in “S4A” using parental cells and a single *FURIN*Δ clone (#2.2.G). The cells were assessed under basal conditions (UT) or after treatment with tunicamycin alone (Tm, 2.5 µg/ml) or in combination with the S1P inhibitor (15 µM) for 6 h. The bar graph shows the mean ± SEM of the normalised median BiP::sfGFP reporter intensity from three independent experiments. All statistical analysis was performed by a two-sided unpaired Welch’s t-test and significance is indicated by asterisks (*p <0.05; ** p < 0.01; *** p < 0.001; ns: non-significant). **(C)** Immunoblot of whole-cell lysates from the indicated genotyped probed with antibodies against FURIN and the loading control actin. FURIN protein levels were determined using HRP or IRDye fluorescently labelled secondary antibodies. On the right, DNA sequence chromatograms of FURIN targeted region are represented, showcasing parental cells and a *FURIN*Δ clone. The rectangle indicates where the FURIN indel mutation occurs, with the sgRNA/Cas9 sequence indicated in a grey rectangle. **(D)** FURIN cleavage assay. *<u>Left panel:</u> FURIN*Δ cells were transfected with a plasmid encoding GFP-ATF6α_TM^S1P,S2P^ ^mut^-LD^WT^ to be used as substrate for *in vitro* FURIN cleavage reactions. 48 h after transfection, cells were lysed, ATF6α was immunoprecipitated (IP) using GFP-Trap Agarose and then subjected to FURIN cleavage by incubating with 1U of commercial FURIN (#P8077S, NEB) for 6 h at 25°C in the presence or absence of calcium (Ca^2+^) Beads were eluted in 2xLB-DTT, boiled for 5 min at 90°C and loaded into 10% SDS-PAGE gels. Membranes were immunoblotted (IB) with an anti-GFP antibody. The expected molecular weight of GFP-ATF6α_TM-^S1P,S2P^ ^mut^-LD^WT^ before cleavage is 67 kDa and 45 kDa post-FURIN cleavage. *<u>Right panel:</u>* As in left panel but using a FURIN control substrate (MBP5-FN-paramyosin-ΔSa). MBP5FN-paramyosin-ΔSa (2.5 µg) was also incubated for 6 h at 25°C in the presence or absence of calcium with 1U of FURIN. The molecular weight of the MBP5-FN-paramyosin-ΔSal substrate before cleavage is 70.6 kDa. FURIN cleavage resulted in an MBP5 mass of 42.8 kDa and a mass of 27.8 kDa. All samples were subjected to SDS-PAGE, and bands were detected by Coomassie blue staining. **(E)** Amino acid-sequence homology between residues 349-670 of human (hs), hamster (cg) and mouse (mo) ATF6α indicating the transmembrane domain (TMD) and the cleavage sites for S1P and S2P proteases. The FURIN putative cleavage site is highlighted in purple. The alignment was performed using the MacVector programme.

**Supplemental Fig S5.**
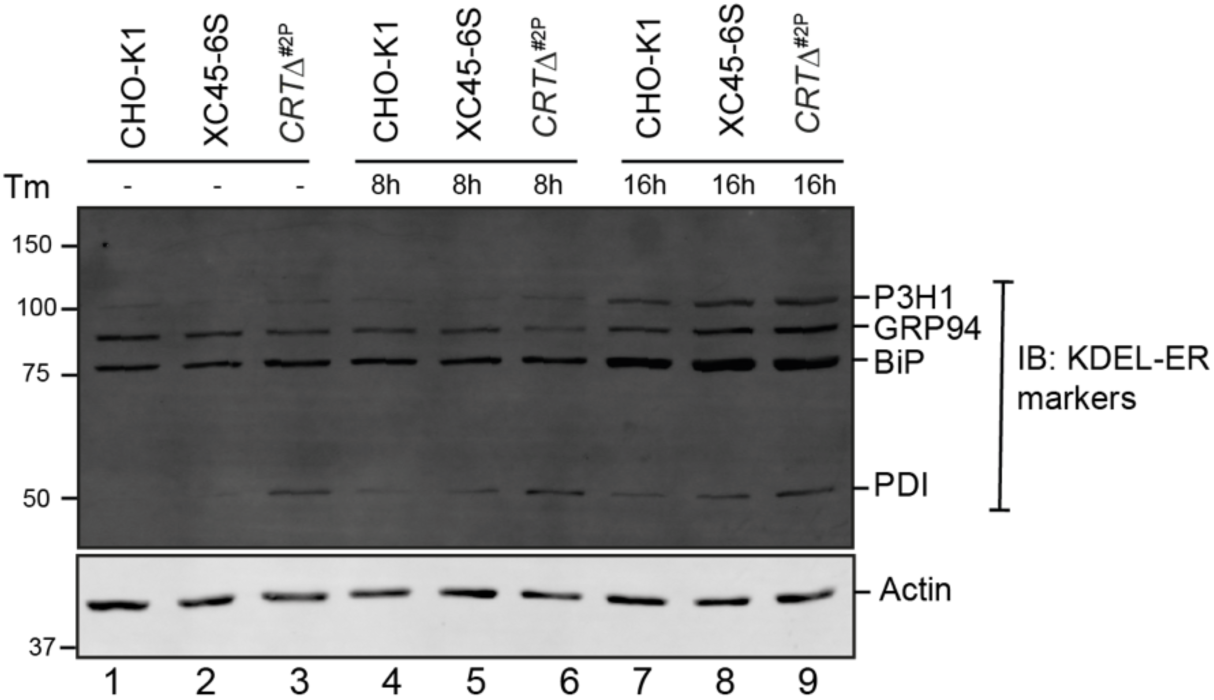
ER-resident proteins (with a KDEL-COOH tag) in CRT depleted cells. Representative immunoblots from two independent experiments show the expression levels of endogenous KDEL receptors in cell lysates from plain CHO-K1 cells, *XC45-6S* dual UPR reporter cells and a derivative *CRT*Δ clone (2P). The samples were also blotted for actin as a loading control.

**Supplemental Fig S6.**
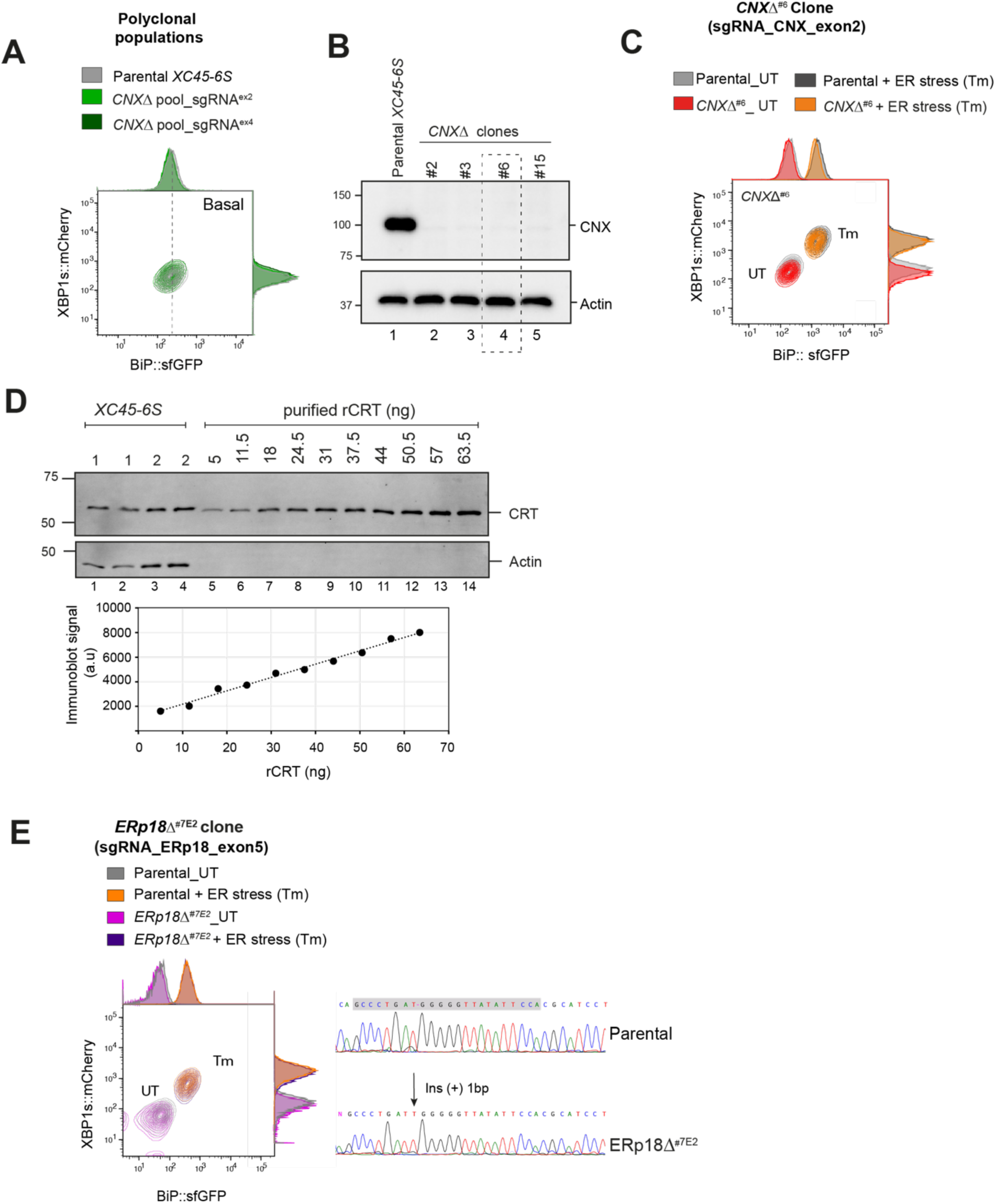
Calnexin depletion does not deregulate ATF6α pathway. **(A)** Two-dimensional contour plots of BiP::sfGFP and XBP1s::mCherry signals in basal conditions in sorted pooled populations where calnexin (CNX) was depleted by CRISPR/Cas9 sgRNAs targeting exon 2 and 4 (in green) and in parental *XC45-6S* cells. **(B)** Immunoblot depicting CNX protein levels in whole-cell lysates from parental cells and four *CNX*Δ derivatives clones. The *CNX*Δ^#6^ clone (highlighted into a rectangle dashed box) was selected for subsequent functional experiments in “S6C”. Actin was used as a loading control marker. **(C)** Contour plot, as in “S6A”, from parental cells and the *CNX*Δ^#6^ single clone treated with Tm for 20 h (black and orange) or left untreated (grey and red). **(D)** Estimation of CRT’s physiological abundance by quantitative immunoblotting. *<u>Top panel:</u>* Immunoblot showing CRT and actin levels for parental *XC45-6S* cells and known amounts of purified bacterially expressed *r*CRT. *<u>Lower panel:</u>* Calibration curve derived from known quantities of purified *r*CRT, fitted to a linear function. Assuming the ER constitutes 10% of cell total volume (Alberts et al., 2002), the concentration of CRT in the ER of CHO-K1 cells was estimated to be around 7 µM. **(E)** *<u>Left panel.</u>* Contour plot, as in “S6C”, from parental cells and the *ERp18*Δ^#7E2^ single clone treated with Tm for 20 h (orange and purple) or left untreated (grey and pink). *<u>Right panel.</u>* DNA sequence chromatograms of ERp18 targeted region, in parental cells and an *ERp18*Δ clone with the arrow indicating the peak where the ERp18 indel mutation occurs. sgRNA Cas9 sequence is indicated with a grey rectangle.

**Supplemental Fig S7.**
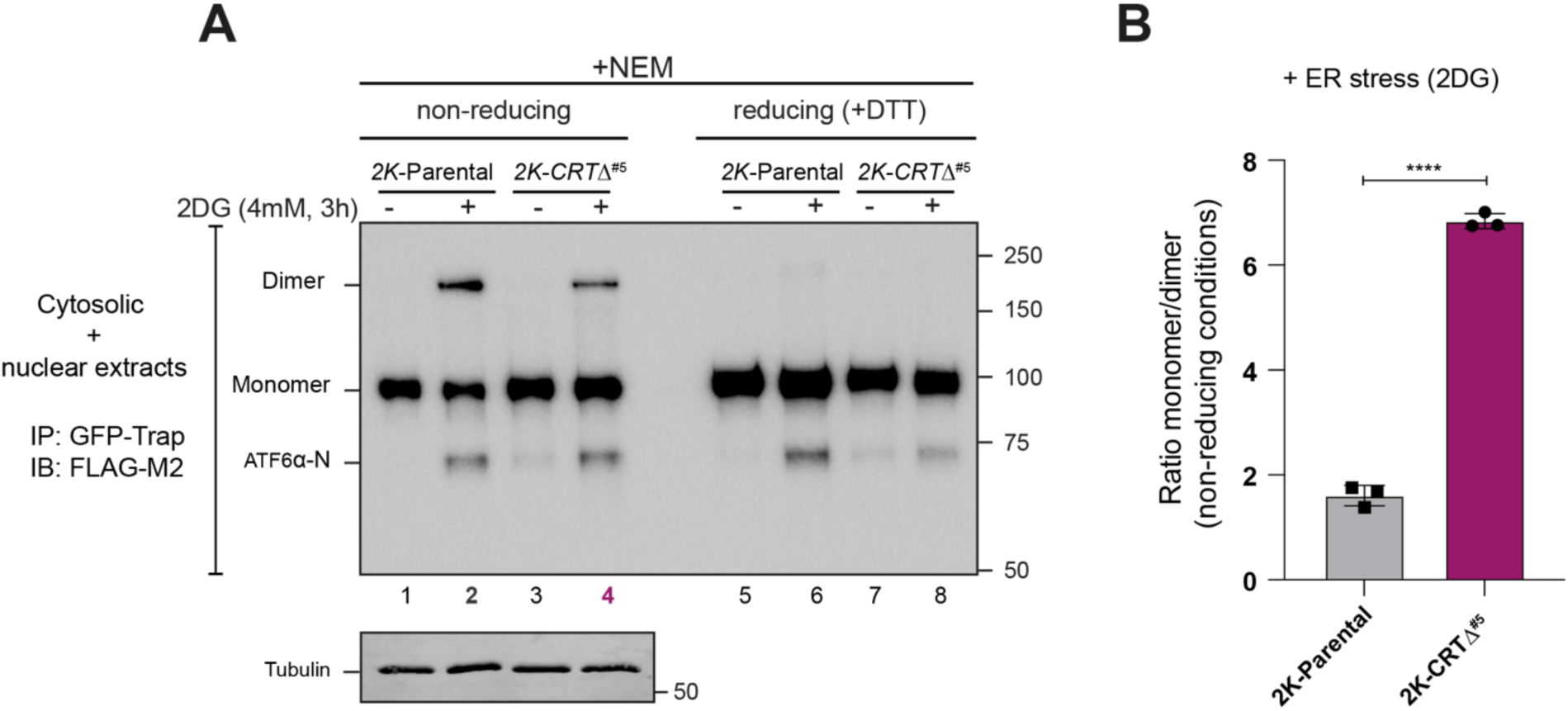
Analyses of ATF6α redox status in CRT-depleted cells. **(A)** Whole-cell lysates from stable *2K* cells with the endogenous *cgATF6*α locus tagged with 3xFLAG and mGreenLantern (mGL) and a derivative *2K-CRT*Δ clone (#5) were separated under non-reducing (-DTT) and reducing (+ 50 mM DTT) SDS/PAGE conditions. ATF6α redox status was analysed under basal conditions and following 2DG treatment (4 mM, 3h) to induce ER stress and ATF6α trafficking. Samples were harvested and lysed in the presence of 20 mM NEM, immunoprecipitated (IP) using GFP-Trap Agarose (ChromoTek) and analysed by immunoblot (IB) using an anti-FLAG M2 antibody to detect ATF6α. Shown is a representative immunoblot from three independent repetitions. **(B)** The graph bar shows the ratio of ATF6α monomer to ATF6α dimer signals in *2K* cells (line 2 from “S7A”) and *2K*-*CRT*Δ clone (line 4 from “S7A”) under stress conditions in non-reducing conditions from three independent experiments indicated as mean ± SEM. Statistical analysis was performed by a two-sided unpaired Welch’s t-test and significance is indicated by asterisks (**** p < 0.0001).

**Supplemental Fig S8.**
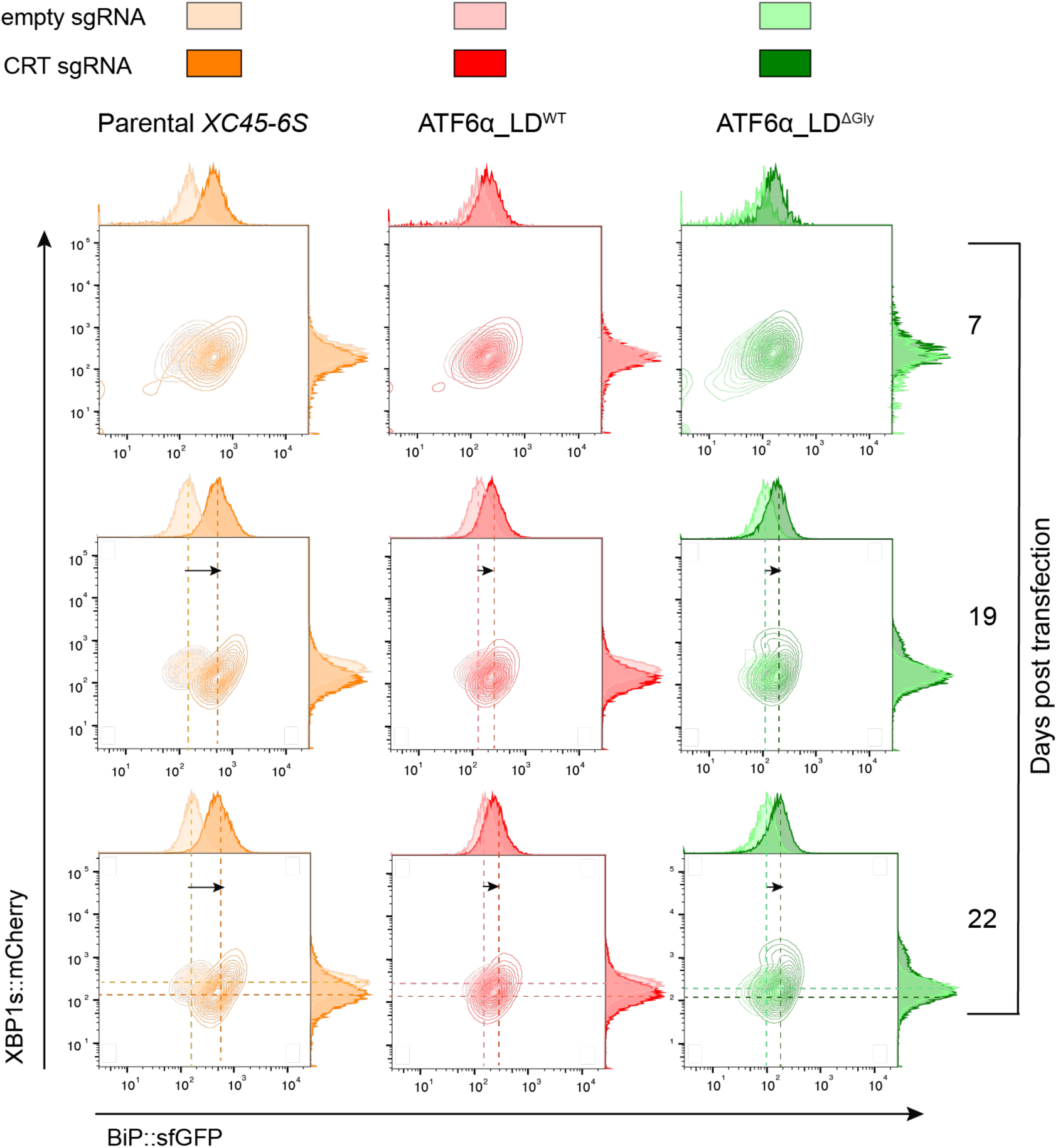
CRT depletion in ATF6α_LD^ΔGly^ knock-in cells does not derepress basally ATF6α reporter activity. Contour plots of BiP::sfGFP and XBP1s::mCherry signals from parental cells (orange) and ATF6α_LD^WT^ (in red) or ATF6α_LD^ΔGly^ (in green) rescued cells (generated in “Fig 5B”) where endogenous CRT was depleted by transient transfection with a sgRNA/Cas9 (Puromycin selection) targeting *cgCRT* locus. Transfected cells were selected in the presence of puromycin (8 μg/ml) and analysed under basal conditions at 7, 19- or 22-days post-transfection. Cells were also transfected with an empty sgRNA/Cas9 plasmid (pale colour) as control for the transfection effect. The presented dataset is representative of a single experiment.

## Supplemental Tables

**Table S1: Ranking of genes enriched in the *activator and repressor ATF6*α *screens*** (see attached file). Genes are ranked by "pos | rank" value. Notably, *MBTPS1*, encoding for S1P protease, is on the top of positively selected genes in the *activator screen,* while *SETDB1* (SET Domain Bifurcated Histone Lysine Methyltransferase 1) is on the top position among the positively selected genes in the *repressor screen*. The top 100 positively selected genes correspond to Fig 1C and Fig 1D.

**Table S2:**
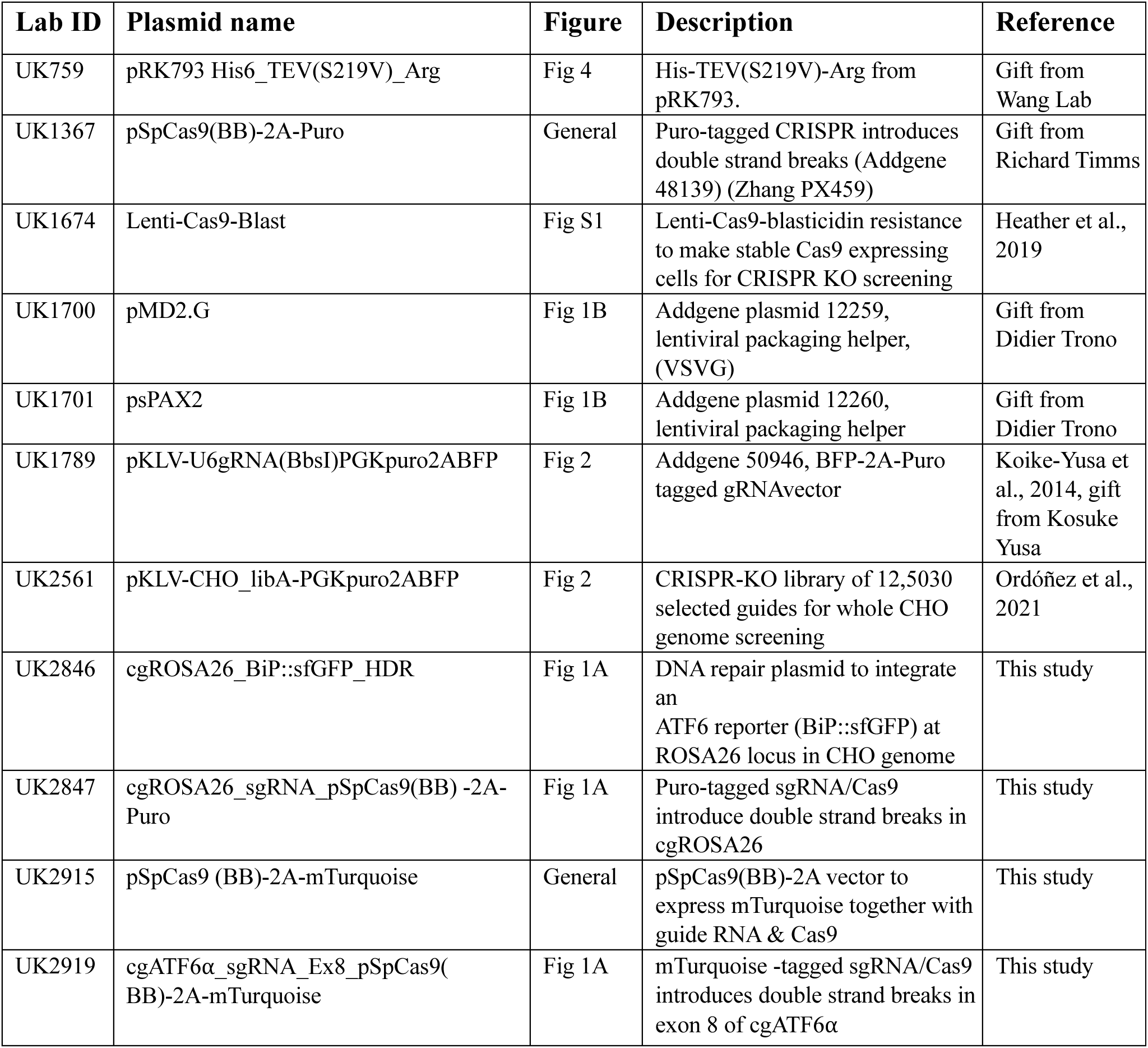
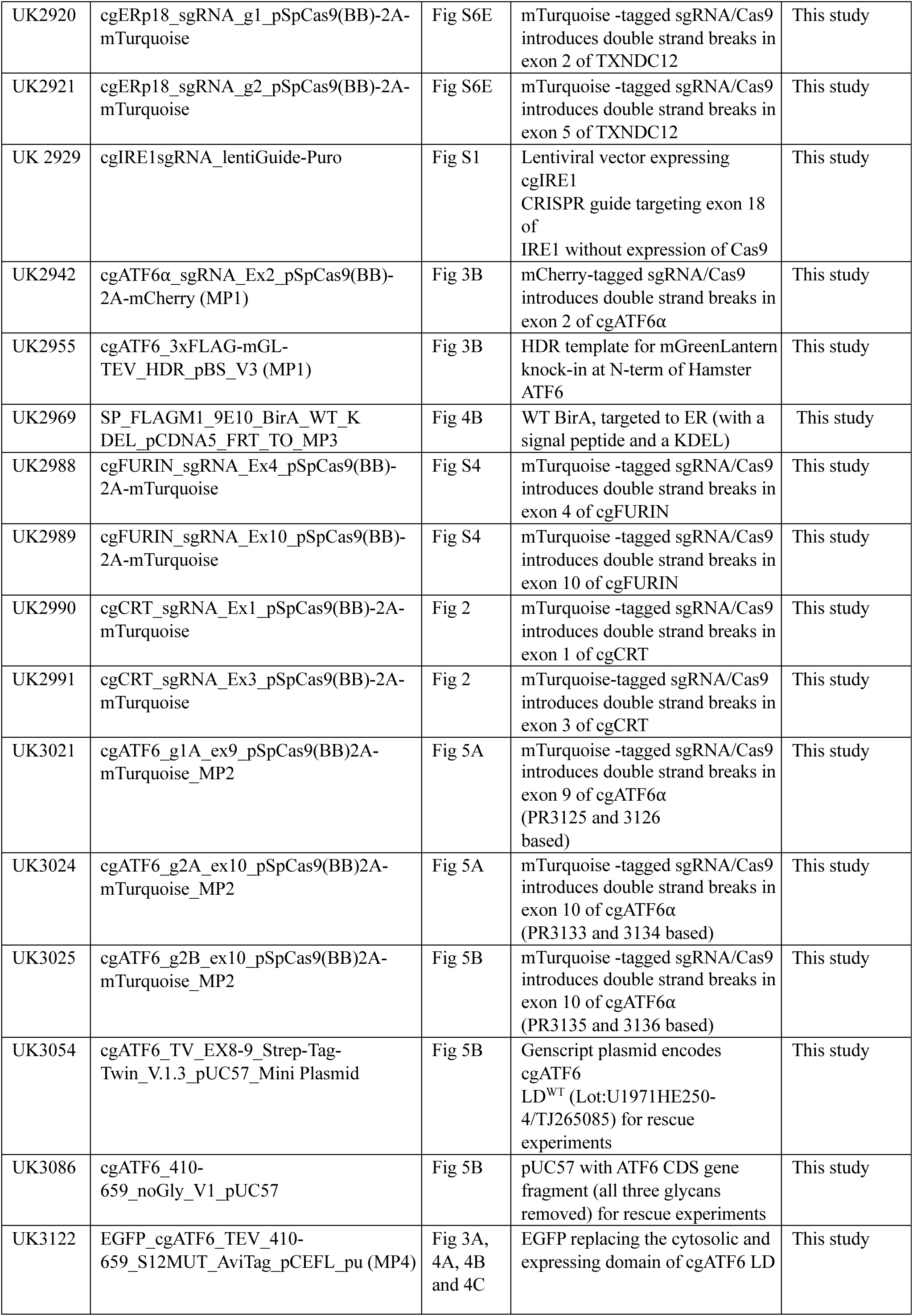
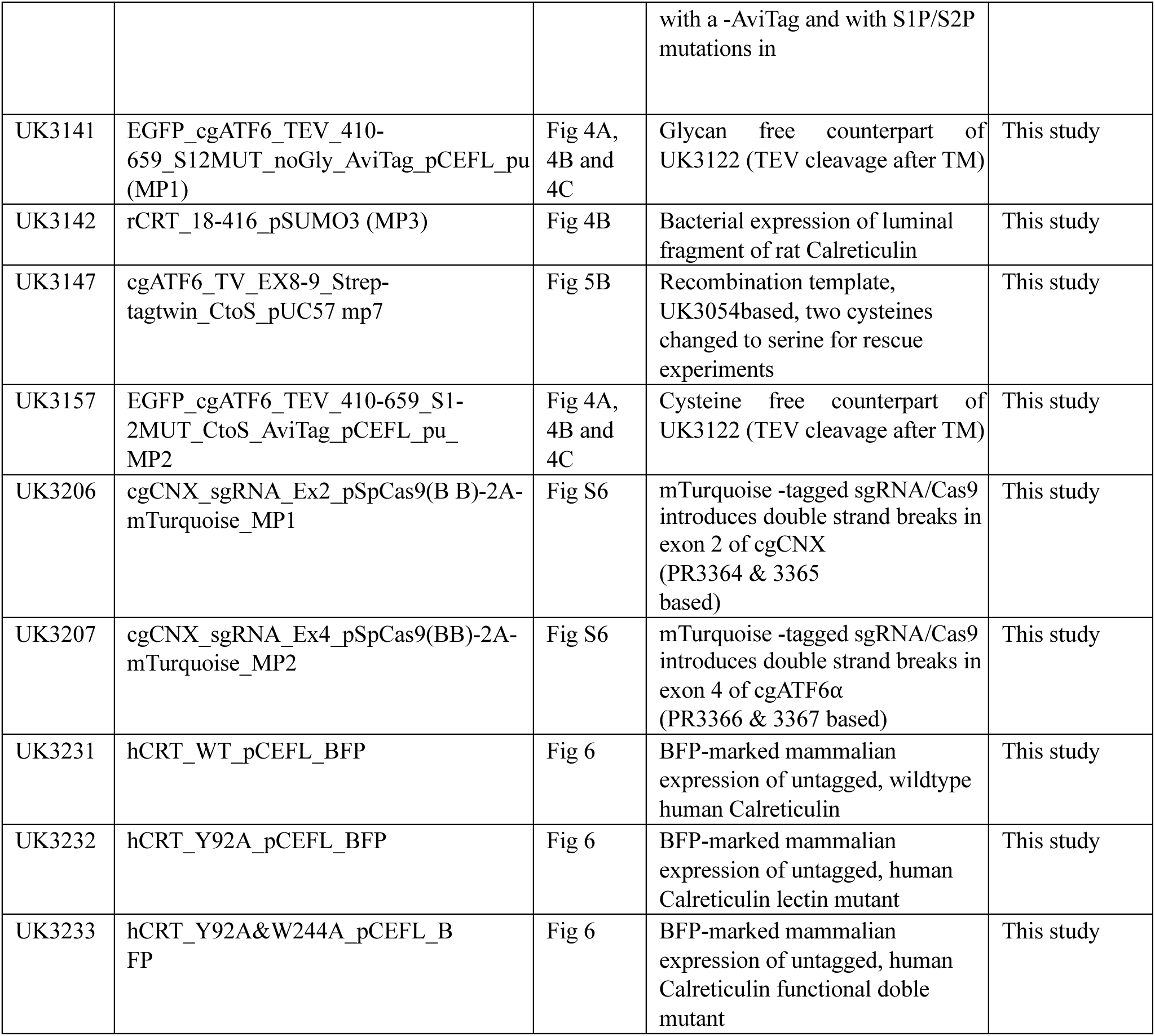
Plasmids used in this study.

**Table S3:**
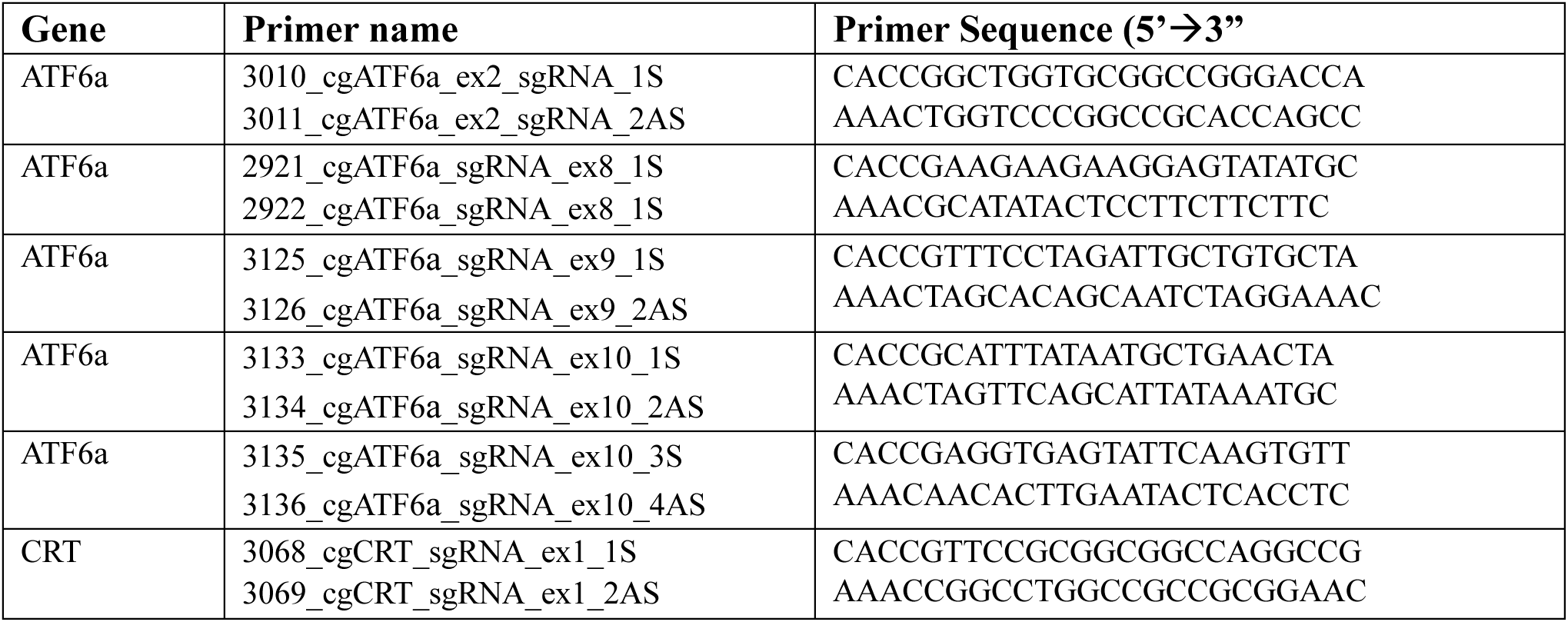
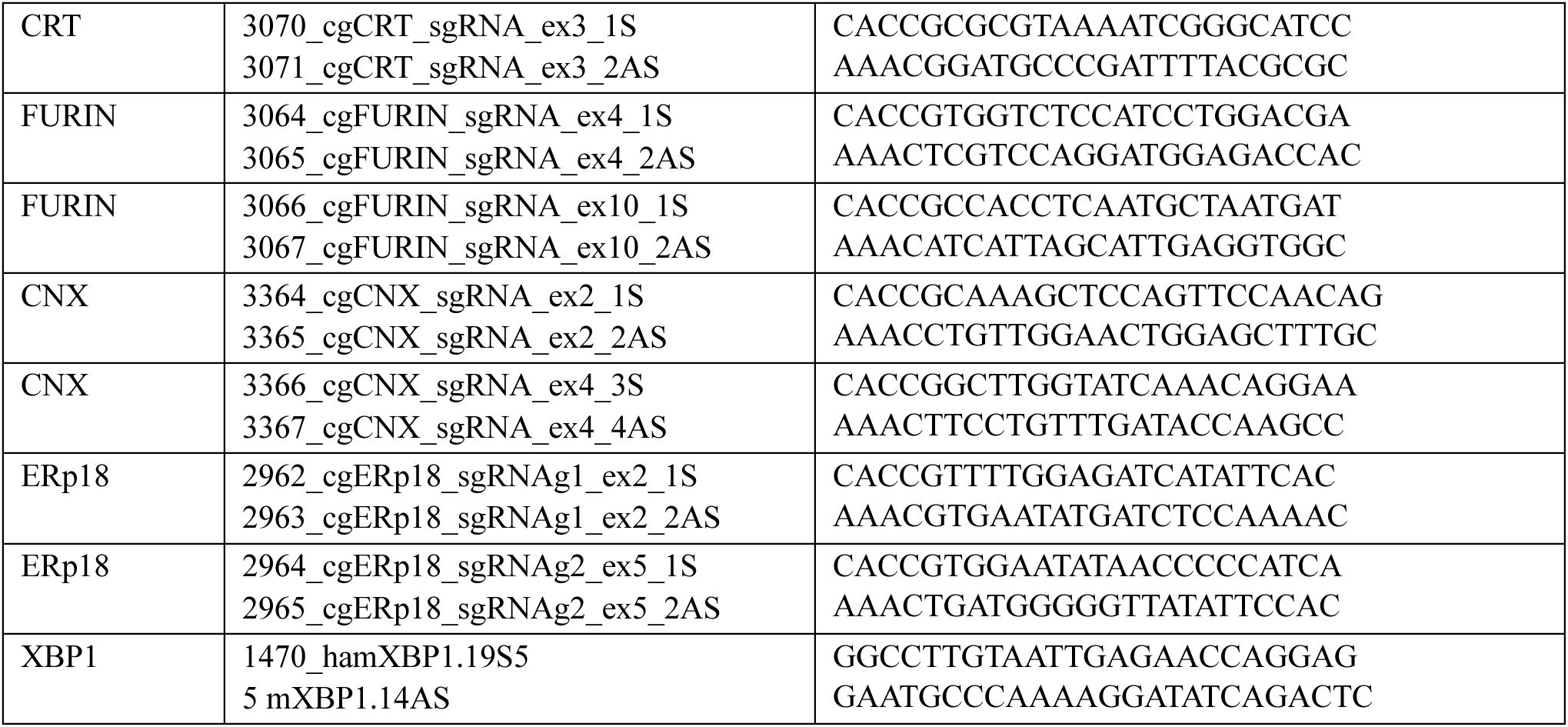
Oligonucleotides used in this study.

**Table S4:**
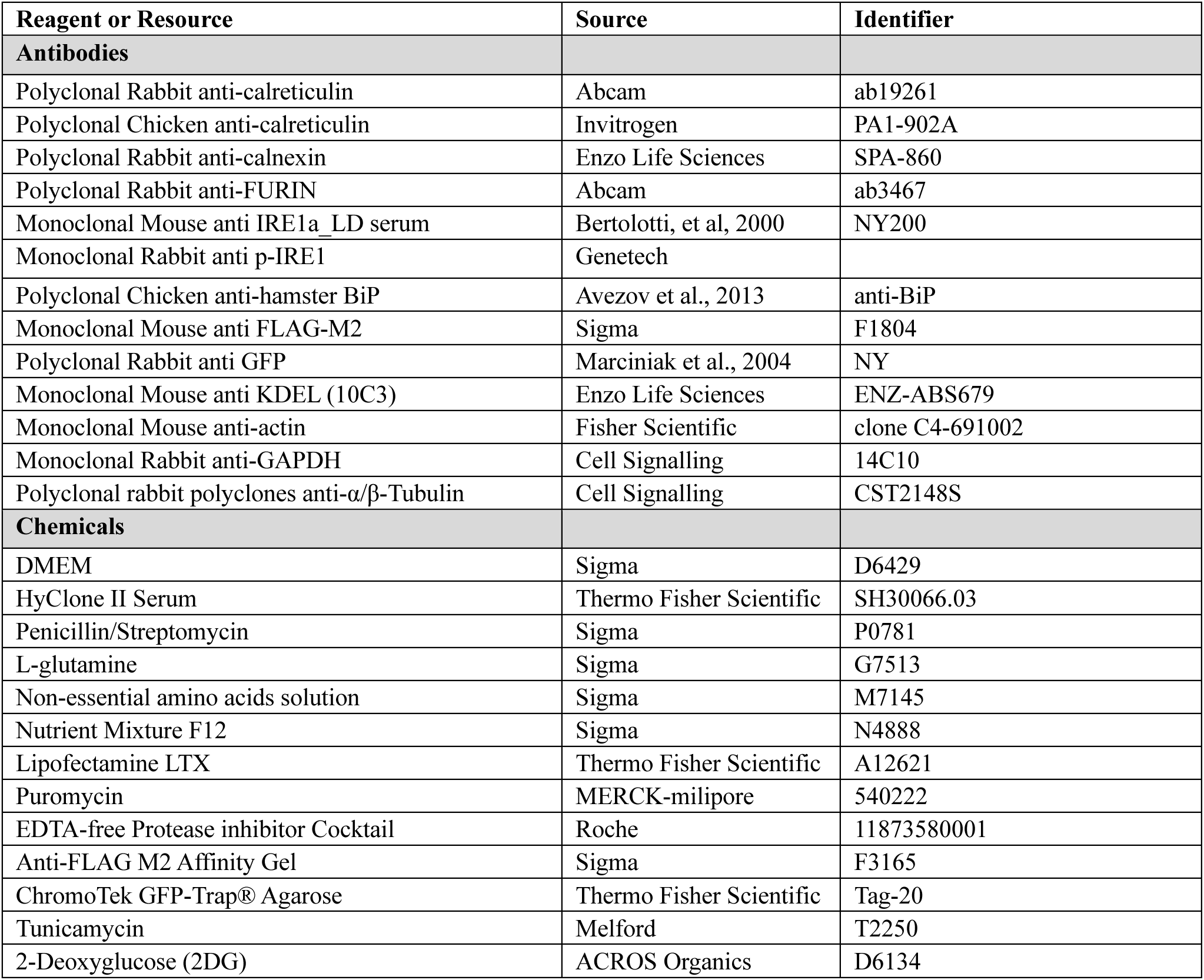
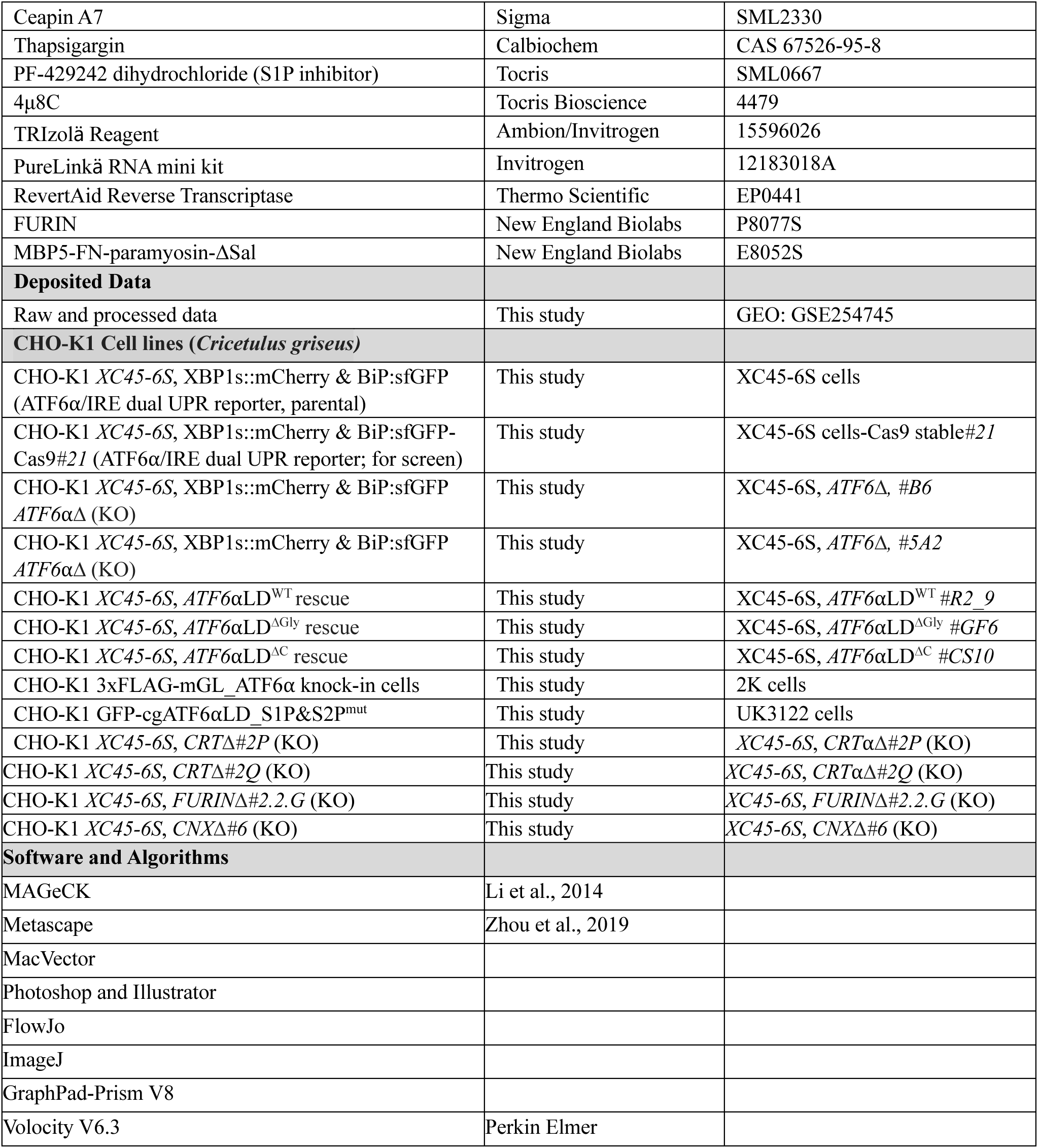
List of antibodies, reagents and software used in this study.

